# Bidirectional interaction between Protocadherin 8 and transcription factor Dbx1 regulates cerebral cortex development

**DOI:** 10.1101/2023.09.28.559903

**Authors:** Andrzej W Cwetsch, Sofia Ferreira, Elodie Delberghe, Javier Gilabert-Juan, Matthieu X. Moreau, Yoann Saillour, Pau Garcia Bolufer, Saray Calvo Parra, Jose González Martínez, Durcia Massoukou, Ugo Borello, Frédéric Causeret, Alessandra Pierani

## Abstract

Brain development requires correct tissue patterning and production of appropriate cell types. Transcription factors (TFs) play essential roles in these processes, regulating the expression of target genes responsible for neuronal subtypes specific features. Cell adhesion molecules are key components of developmental processes that control cell sorting, migration, neurite outgrowth/guidance and synaptogenesis. To date, the link between TFs and cell adhesion molecules is considered to be unidirectional. Here, we demonstrate that ectopic expression of Dbx1 leads to spatio-temporally restricted increased expression of *Pcdh8* and cell aggregation, together with changes in neuronal identity. Surprisingly, Pcdh8 overexpression also induces Dbx1 expression as well as a complete reorganisation of apico-basal polarity and dorso-ventral patterning *via* Notch signalling. Altogether, our work therefore points to cell adhesion molecules as unexpected, yet important, players in the regulation of cell identity and, in particular, Pcdh8 through its bidirectional interaction with the Dbx1 transcription factor.

## INTRODUCTION

Early steps of brain development rely on tightly controlled action of transcription factors (TFs) that organize the dorso-ventral (DV) axis of the central nervous system (CNS). The precise expression of TFs in domains along the DV axis ensures robust patterning and their faulty regulation leads to the disruption of the DV organisation and/or cell identity (Arai et al., 2019; Godbole et al., 2017; Leung et al., 2022; Marklund et al., 2010; Pierani et al., 1999). Among other TFs, the developing brain homeobox 1 (Dbx1) was shown to function as a potent DV patterning and cell-fate determinant in different regions of the CNS, such as the mouse spinal cord (Pierani et al., 2001), rhombencephalon (Bouvier et al., 2010a) and diencephalon (Sokolowski et al., 2016). In the mouse developing telencephalon, the Dbx1 gene exhibits a very precise expression in ventral pallium (VP) progenitors just above the pallial-subpallial boundary (PSB) and the septum (Bielle et al., 2005). Recently, it was shown that its ectopic expression in the dorsal developing cerebral cortex (Arai et al., 2019) promotes Cajal-Retzius (CRs) and SP-like cells fate (Arai et al., 2019), with SP fate being dependent on primate-specific *cis*-acting elements acting in postmitotic neurons. Furthermore, in the spinal cord, Dbx1 has been shown to regulate DV patterning expression profiles of TFs and to establish subtype-specific interneuron fate (Pierani et al., 2001). This is mediated by controlling the DV expression profile of Notch ligands, namely suppressing the expression of Jag1 and promoting that of Delta1 (Marklund et al., 2010). Last, Dbx1 was reported to drive cell cycle exit in birds and mice (Arai et al., 2019; García-Moreno et al., 2018; Nomura et al., 2008). However, it remains unknown how Dbx1 influences cell fate at the molecular level in the developing cerebral cortex (Leung et al., 2022).

Protocadherins (Pcdhs) are cell adhesion molecules belonging to the cadherin superfamily and are classified into two subgroups in vertebrates: clustered (cPcdhs) and non-clustered Pcdhs (ncPcdhs). Their expression is regulated in a spatiotemporally manner during development, being involved in multiple processes required for proper CNS formation. In particular, δ-Pcdhs (i.e. Pcdh8, Pcdh9 and Pcdh19), a subgroup of ncPcdhs, have been associated with dendrite and axon formation, morphology, migration and guidance (Bassani et al., 2018; Biswas et al., 2010; Bruining et al., 2015; Cooper et al., 2015; Cwetsch et al., 2022; Pancho et al., 2020; Yasuda et al., 2007). Moreover, human mutations in some δ-Pcdhs are associated with several neurodevelopmental disorders, such as autism, schizophrenia and epilepsy (Kahr et al., 2013). Additionally, in the mammalian neocortex, it was shown that patterned Pcdh expression biases the fine spatial and functional organization of individual neocortical excitatory neurons (Lv et al., 2022). However, whether Pcdhs play a role in regulating TF expression to establish neuronal identities or DV patterning in the embryonic murine brain remains unknown.

We previously demonstrated that at embryonic day 11.5 (E11.5), ectopic expression (EE) of Dbx1 (hereafter Dbx1 EE) in the dorsal pallium (DP) of the developing mouse brain imparts cell identity by inducing ectopic Cajal-Retzius (CRs) and subplate (SP)-like neuron fate (Arai et al., 2019), thus resulting in changes in cell fate in the DP. The molecular mechanism behind this process, however, remains unknown. Here, we report that Dbx1 EE induces cell aggregation through Pcdh8 upregulation and that Pcdh8 is required for Dbx1-driven neuronal fate specification, particularly the generation of Nr4a2^+^ claustro-amygdalar complex neurons (CLA). We used single-cell RNA sequencing (scRNAseq) to characterize distinct expression dynamics for both genes during neuronal differentiation at the VP and septum. Consistently, we reveal that Pcdh8 EE induces a cell-autonomous (CA) upregulation of Dbx1 as well as non-cell-autonomous (NCA) defects in cell proliferation and tissue patterning, resulting in a massive reorganization of the cortical primordium. Finally, we identify the Notch ligand Jag1 as a downstream effector of Pcdh8. Our findings therefore point to cell adhesion molecules, in particular Pcdh8, as players acting not only downstream, but also upstream of TFs during brain development.

## RESULTS

### Ectopic Dbx1 expression in the lateral pallium induces cell aggregation

Using *in utero* electroporation (IUE), we previously investigated the effects of ectopic pCAGGS-Dbx1-iresGFP expression in the E11.5 mouse DP. We found that Dbx1 EE induced the production of neurons harbouring features of Cajal-Retzius (Reln^+^/Calr^+^ (Calretinin/Calb2)) and SP-like Nr4a2^+^Bcl11b^+^ (Nurr1^+^Ctip2^+^) fates 48h after electroporation (Arai et al., 2019). Surprisingly, when electroporation of Dbx1 at E11.5 encompassed lateral and ventral pallial domains, we noticed 48h later, that transfected cells tended to aggregate forming cohesive clusters (Figure 1A, B, Figure S1A, C). Such clusters were preferentially formed in the lateral pallium (LP) (Figure 1C, D) at the border between the ventricular/subventricular zone (VZ/SVZ) and the cortical plate (CP) (Figure 1B, Figure S1C). They were rarely observed in the dorsal or medial cortex, and in GFP-electroporated control brains (Figure 1B, D and data not shown). Aggregation also failed when IUE was performed one day later, at E12.5, suggesting a spatio-temporal modulation in the response to Dbx1 EE (Figure S1A, B). To assess cell adhesiveness differences following Dbx1 EE, we performed an adhesion assay, as previously described (Porlan et al., 2014). Neuronal stem cells were transfected *in vitro* with Dbx1 EE (iresGFP) or a control vector (iresGFP) and after 24h, GFP^+^ cells were seeded onto NC929 cell monolayers (overexpressing Ncad/Cdh2), and the number of attached cells was quantified (Figure S1E). The number of attached GFP^+^ cells, normalized to transfection efficiency (total GFP^+^ cells before seeding quantified by fluorescence-activated cell sorting (FACS)), showed that Dbx1 EE significantly enhanced adhesion properties (Figure S1F, G). We then investigated whether Dbx1 EE and increased adhesion influenced cell fate. Consistent with our previous findings, we found numerous cells expressing Nr4a2 within the Dbx1-induced aggregates, but not in control GFP-electroporated cells (Figure 1B, E, G and Figure S1H). However, these clusters lacked expression of Ctip2/Bcl11b, a marker of early generated neurons and SP neurons, unlike what is observed following IUE of the DP (Arai et al., 2019). Instead, Ctip2 appeared excluded from the GFP^+^ regions (Figure 1F, H and Figure S1I). Indeed, when we measured immunofluorescence (IF) intensity of Nr4a2^+^ or Ctip2^+^ neurons aligned with the GFP signal of Dbx1 EE cells, we observed a positive correlation with Nr4a2 but a negative correlation with Ctip2 (Figure 1G, H). Surprisingly, Dbx1-induced aggregates were also Reln^−^ and Calb2^−^, with some individual Reln^+^ and Calb2^+^ cells at the cortical surface (Figure S1J, K). Additionally, the deep layer marker Tle4 was not detected in Dbx1 EE-expressing cells (data not shown). All electroporated GFP^+^ cells expressed Dbx1 (Figure S1C, D). When embryos were collected 7 days after IUE (E11.5-18.5), Dbx1 EE^+^/Nr4a2^+^ cells accumulated in the intermediate zone (IZ)/SP (Figure 1I). Within the CP, Dbx1 EE^+^ were Nr4a2^−^ and coexisted with Nr4a2^+^ non-electroporated cells (Figure 1I), the latter being endogenous late-born glutamatergic neurons of the lateral cortex (Fang et al., 2021). To determine whether Nr4a2^+^ neurons derive from Dbx1-expressing progenitors in the developing cerebral cortex, we used *Dbx1*^*nlsLacZ/+*^ mice (in which the LacZ gene coding for β-galactosidase (β-gal) reporter is inserted directly under the control of the *Dbx1* promoter), allowing short-term lineage tracing of Dbx1-derived cells. We confirmed that β-gal^+^ Dbx1-derived cells located in the postmitotic compartment of the VP expressed Nr4a2 at both E12.5 and E14.5 (Figure S2A and data not shown). Furthermore, we used *Dbx1*^*Cre*^*;Rosa26*^*YFP*^ animals to permanently trace Dbx1-derived progeny. In the rostral LP at E12.5 and later at E14.5, we did not detect YFP^+^Nr4a2^+^ in the SP (Figure S2B, C), consistent with the previously reported at E16.5 (Arai et al., 2019). This shows that Dbx1 progenitors do not give rise to Nr4a2^+^ SP neurons in the mouse. Instead, in intermediate RC sections at the VP, we observed ventrally located groups of YFP^+^Nr4a2^+^ cells at both at E12.5 and E14.5 (Figure S2B, C, panel c). Together with data showing *Nr4a2*^+^ Dbx1-derived neurons in a E14.5 dataset using *Dbx1*^*Cre*^*;Tau*^*LacZ*^ reporter (data not shown), these findings indicate that Dbx1 progenitors in the mouse VP originate pallial Nr4a2^+^ neurons that migrate ventrally, corresponding to neurons of the future claustro-amygdaloid complex (CLA) (Puelles et al., 2016a), and not SP neurons.

**Figure 1.**
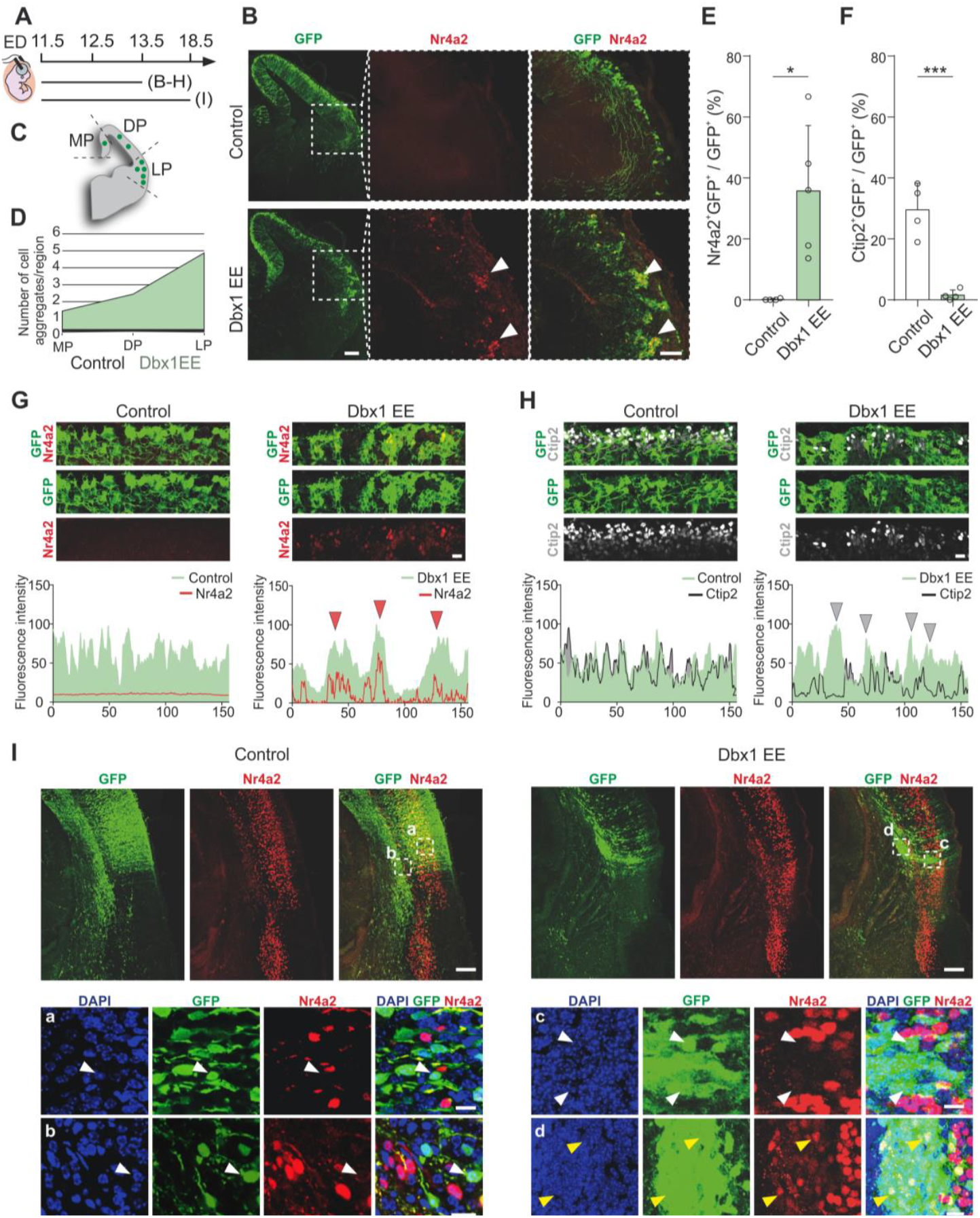
Dbx1 EE induces cell aggregation in the lateral pallium. (A) Schematic timeline of the IUE. ED, electroporation day. (B) Confocal images of GFP (green) in coronal sections of E13.5 mouse brain cortices electroporated at E11.5 with control GFP or Dbx1 EE, co-labeled with Nr4a2 (red). Dashed squares magnified on the right. White arrowheads indicate aggregates. Scale bars: 200 μm, 100 μm (magnified). (C) Aggregate analysis scheme in MP-medial, DP-dorsal, LP-lateral pallium. (D) Aggregate formation analysis in control (n=6) and Dbx1 EE (n=6) sections as showed in (C). (E, F) Percentages of (E) Nr4a2+GFP+ and (F) Ctip2+GFP+ cells among GFP+ cells. Data are mean ± SEM; circles represent values from independent electroporations (n=5 (E); n=4 (F)); Student’s t-test, (E) *p=0.01, (F) ***p=0.0008. (G, H) Top: confocal images of E13.5 mouse coronal sections showing GFP+ (green) cells electroporated with control or Dbx1 EE, co-labeled with (G) Nr4a2 (red) or (H) Ctip2 (gray). Scale bar: 25 μm. Bottom: fluorescent intensities of GFP+ cells (green area) and Nr4a2+ (red line, G) or Ctip2+ (gray line, H), along the cortical surface (μm). Red (G) or gray (H) arrowheads show positive (Dbx1 EE GFP+ areas and Nr4a2 labeling) or negative (Dbx1 EE GFP+ areas and Ctip2 labeling) fluorescent intensity correlation. (I) Top: confocal images of GFP (green) in coronal sections of E18.5 mouse brain cortices electroporated at E11.5 with control vector or Dbx1 EE, co-labeled with Nr4a2 (red) and DAPI (blue) counterstaining. Bottom: Dashed squares magnified (a, b, c, d). White and yellow arrowheads indicate Dbx1 EE-/Nr4a2+ and Dbx1 EE+/Nr4a2+ cells in the electroporated area. Scale bars: 200 μm, 25 μm (magnified). See also Figures S1 and S2.

Altogether, these results show that Dbx1 EE promotes cell-cell adhesion in a CA manner at the LP and VP and this gives rise to Nr4a2^+^ CLA neurons. In addition, we found that Dbx1-induced cell aggregation is spatio-temporally restricted by the competence of the cortical neuroepithelium with no aggregation in DP, correlating with Dbx1 EE promoting exclusively CR and SP-like fates at E11.5 (Arai et al., 2019), and aggregation in LP/VP with induction of CLA neurons. Moreover, we confirm that endogenous VP Dbx1^+^ progenitors do not generate SP neurons in mice, and SP-like fate arises from primate-like Dbx1 expression in postmitotic neurons (Arai et al., 2019), whereas CR and CLA neurons are endogenous fates driven by VP Dbx1^+^ progenitors in mice.

### Pcdh8 and Dbx1 show concomitant expression in pallial progenitors and Dbx1-derived neurons

To begin investigating the molecular mechanisms behind Dbx1-induced aggregation, we took advantage of previously published bulk transcriptomic profiles of FACS-purified Dbx1-derived cells from dorsomedial and dorsolateral cortices of E12.5 *Dbx1*^*Cre*^*;Rosa26*^*YFP*^ embryos (Figure 2A, B, C) (Griveau et al., 2010). We compared the expression level of genes belonging to the Protocadherin (Figure 2B) and Cadherin (Figure 2C) families, two major players in neuronal adhesion (Halbleib and Nelson, 2006; Peek et al., 2017; Suzuki and Takeichi, 2008). We found that *Cdh2* (also known as *Ncad*), *Pcdh8, Pcdh9* and *Pcdh19* stood out with ~10-fold higher expression levels in comparison to other adhesion molecules (Figure 2B, C). To better appreciate their possible implication downstream of Dbx1, we used scRNAseq. We first used a previously generated dataset sampling cell diversity around the pallial-subpallial boundary (PSB) at E12.5 (Moreau et al., 2021). We found *Cdh2, Pcdh9* and *Pcdh19* broadly expressed, with no specific enrichment (https://apps.institutimagine.org/mouse_pallium/), which was confirmed by *in situ* hybridization (ISH) (Supplementary Figure 5G-I). In contrast, *Pcdh8* displayed an interesting expression pattern, with strong mRNA enrichment in progenitor domains adjacent to the most characteristic telencephalic regions of Dbx1 expression, specifically the VP dorsal to the PSB and the septum, on embryonic *wild-type* (WT) brain sections (Bielle et al., 2005) (Figure 2G, Figure S3C and Figure S5G-I). Particularly, we detected *Pcdh8* expression in apical progenitors (APs) of the VP, subpallium and septum (Figure 2D, F and Figure S3A), but not of LP and DP regions. Since *Dbx1* expression is initiated in APs of the VP and strongly increases during the transition to the basal progenitor (BP) state (Figure 2D), we compared the temporal dynamics of both *Dbx1* and *Pcdh8*. First, we used a previously reconstructed differentiation trajectory, in which VP lineage cells are ordered along a pseudotime axis representing their differentiation state, from APs to neurons at E12.5 (Moreau et al. 2021) (Figure 2E). We found that *Pcdh8* and *Dbx1* are expressed sequentially in the VP lineage, with *Pcdh8* peaking in APs, followed by *Dbx1* in BPs and *Nr4a2* in neurons (Figure 2E). To further investigate the co-expression of *Dbx1* and *Pcdh8* at earlier stages in the VP, we produced a scRNAseq dataset of the entire telencephalic vesicle at E11.5-E12 and detected *Dbx1*^*+*^*Pcdh8*^*+*^ cells at the AP-to-BP transition (Figure 2F), suggesting a possible, albeit transient, direct interaction between Dbx1 and Pcdh8. We also performed scRNAseq on septal explants collected from E11.5 *PGK*^*Cre*^*;Rosa26*^*YFP*^ and E12.5 *Dbx1*^*Cre*^*;Rosa26*^*Tomato*^ embryos, the latter enabling genetic tracing of Dbx1-derived cells (Figure S3A, B) (https://apps.institutimagine.org/mouse_septum/). We observed a robust co-expression of *Dbx1* and *Pcdh8* in both progenitors and septal-derived postmitotic excitatory neurons (*Tbr1*^−^/*Slc17a6*^+^) (Figure S3A, B). We further validated the co-expression of *Dbx1* and *Pcdh8* by fluorescent ISH (FISH) at E12.5 (Figure 2H) by observing co-expressing cells within the postmitotic compartment of the septum (Figure 2H panels 1a, b) in a ventrally migrating stream of postmitotic Dbx1-derived cells (Bielle et al., 2005; Griveau et al., 2010) (Figure 2H panel 2c). Additionally, we detected co-expression in progenitors at the VP/PSB (Figure 2H panel 3d). At the protein level, Pcdh8 expression in the VP and in Dbx1-derived neurons was confirmed by co-immunostaining of β-gal and Pcdh8 (antibody validation at E12.5, Figure 2G) on brain sections of E11.5 and E12.5 *Dbx1*^*LacZ/+*^ embryos (Figure S4A, B). In the postmitotic compartment of the VP of E12.5 WT animals, we found Pcdh8^+^Nr4a2^+^ co-expressing neurons (Figure S4C). Finally, to confirm the epistasis between Dbx1 and Pcdh8, we analysed *Pcdh8* expression in E11.5 *Dbx1*^*LacZ/LacZ*^ knockout (KO) animals and observed changes in *Pcdh8* expression domains at the septum and VP/PSB as well as in the localization of *Pcdh8*^*+*^ cells in ventral and lateral postmitotic compartments (Figure S3C).

**Figure 2.**
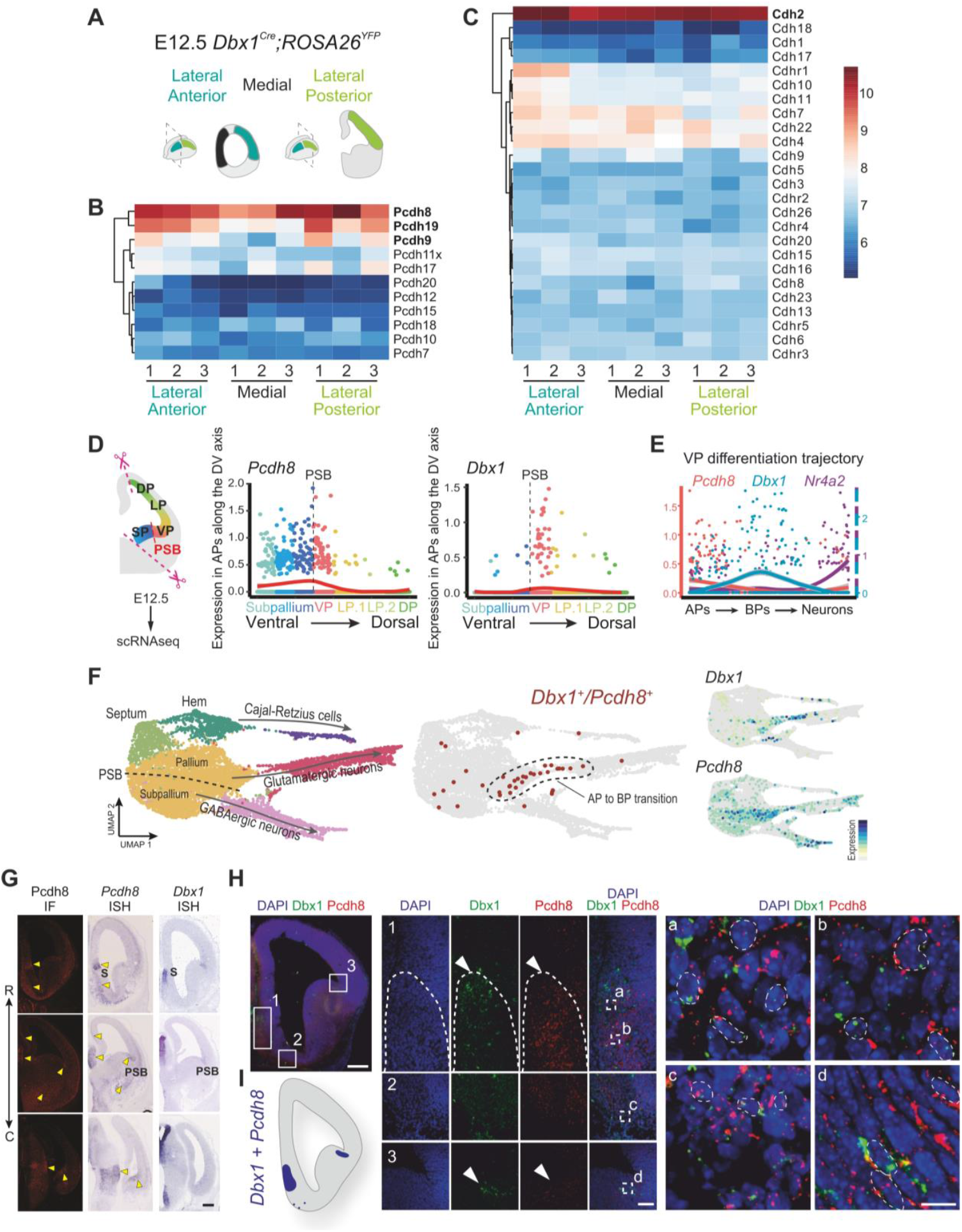
The endogenous expression of Pcdh8 and Dbx1 partially overlaps. Experimental design: E12.5 *Dbx1*^*Cre*^*;Rosa26*^*YFP*^ brain with marked dissected regions of lateral anterior, medial and lateral posterior areas used for bulk transcription profiling of FACS-sorted YFP^+^ cells. **(B, C)** Heatmaps of microarray analysis from E12.5 *Dbx1*^*Cre*^*;Rosa26*^*YFP*^ brains (*n*=3) showing expressing levels of **(B)** protocadherins (Pcdhs) and **(C)** cadherins (Cdhs). The color scale (right) represents expression levels from low (blue) to high (red). **(D)** Left: Scheme of the dissected brain region from E12.5 *wild-type* (WT) subjected to scRNAseq. Right: Comparison of *Pcdh8* and *Dbx1* gene expression trends along pseudo dorso-ventral (DV) scores. Each dot represents a single apical progenitor (AP), color-coded by domain: SP, subpallium; VP, ventral pallium; LP, lateral pallium; DP, dorsal pallium; PSB, pallial-subpallial boundary. The red curve indicates the smoothed expression profile. (E) Comparison of Pcdh8, Dbx1 and Nr4a2 gene expression trends along the VP differentiation trajectory, from APs to basal progenitors (BPs) to neurons, mapped onto a pseudotime axis. **(F)** Left: UMAP representation of the scRNAseq dataset from the entire telencephalic vesicle at E11.5-E12 WT brains, with cells color-coded by cell type. Neuron types are indicated, and arrows represent the main differentiation trajectories. The dashed line marks the PSB. Cells co-expressing *Dbx1* and *Pcdh8* (dark red) are mostly found at the transition from AP to basal progenitor (BP), as outlined by the dashed area. Right: UMAP visualization of *Dbx1* and *Pcdh8* gene expression. **(G)** Left: Confocal images of Pcdh8 (red) immunofluorescence (IF) in coronal sections of E12.5 WT mouse brain. Right: Bright-field images of *Pcdh8* and *Dbx1 in situ* hybridization (ISH) in coronal sections of E12.5 WT mouse brains. The double arrow line indicates the rostro-caudal (R-C) axis. Yellow arrowheads indicate strong correspondence between Pcdh8 signals in IF and ISH, validating the anti-Pcdh8 serum. S, septum. Scale bar: 100 μm. **(H)** Confocal images of WT E12.5 mouse brain coronal sections showing fluorescence ISH (FISH) of *Dbx1* (green) and *Pcdh8* (red), co-stained with DAPI (blue). Selected areas 1 to 3 represent: (1) medio-ventral part of the septum postmitotic compartment; (2) postmitotic compartment of pallial cells migrating ventrally; (3) PSB. White arrows indicate areas with the highest Dbx1 and Pcdh8 expression. Dashed squares (a, b, c, d) magnified on the right panel with dashed circles outlining cells co-expressing *Dbx1* and *Pcdh8*. **(I)** Schematic representation of overlapping regions with high *Dbx1* and *Pcdh8* expression detected by FISH. Scale bars: 100 μm (upper left panel), 25 μm (1, 2, 3), 5 μm (a-d). See also Figures S3, S4 and S5.

We therefore conclude that *Dbx1* and *Pcdh8* display interrelated dynamic expression patterns in the developing telencephalon, occurring both concomitantly in AP/BP progenitors between E11.5 and E12.5, and sequentially during neuronal differentiation in both the VP and septal lineages, suggesting possible direct interaction. Furthermore, co-expression studies combined with tracing analysis confirm the existence of endogenous Dbx1^+^Pcdh8^+^ and Dbx1-derived Nr4a2^+^Pcdh8^+^ progenitors and neurons in the embryonic telencephalon.

### Dbx1 EE induces Pcdh8 expression

To address whether Dbx1 EE regulates *Pcdh8* expression, we performed quantitative reverse transcription polymerase chain reaction (RT-qPCR) on lateral cortical tissue dissected 48h post-electroporation (E11.5-13.5) with Dbx1-iresGFP or GFP control expression vectors (Figure 3A, B). We included in the analysis *Reln, Calb2* and *Nr4a2* given their known upregulation upon Dbx1 EE in the DP (Arai et al., 2019), *Ctip2* given its exclusion from Dbx1-induced aggregates (Figure 1F, H and Figure S1I), and *Pcdh9, Pcdh19* and *Ncad*. As expected, *Dbx1* mRNA levels increased in Dbx1-electroporated regions compared to controls (Figure 3C). The expression levels of *Pcdh8, Nr4a2* and *Calb2* transcripts were significantly increased in the electroporated areas, while the remaining genes remained unchanged (Figure 3D). Similar results were obtained 24h post-electroporation (E11.5-12.5), with a significant upregulation of *Dbx1, Pcdh8* and *Nr4a2* transcripts, arguing for a rapid, if not direct, regulatory effect (Figure 3E, F). Additionally, following Dbx1 EE (E11.5-13.5), we observed a significant negative correlation between *Dbx1* and *Pcdh19* gene expression (Figure S5A), which was not the case for other cadherins (i.e. *Pcdh8, Pcdh9* and *Ncad*) (Figure S5A), pointing to other possible partners in gene co-regulation networks. This correlated with *Pcdh19* expression on sections in DP but not LP/VP progenitors (Figure S5H-I). By contrast, Dbx1 IUE at E12.5 (E12.5-14.5) no longer altered *Pcdh8* and *Nr4a2* expression, but resulted in the upregulation of *Pcdh9, Pcdh19, Ncad*, and *Ctip2* (Figure S5B-E), which overall aligns with the absence of cell aggregates upon Dbx1 EE at this developmental stage (Figure S5F).

**Figure 3.**
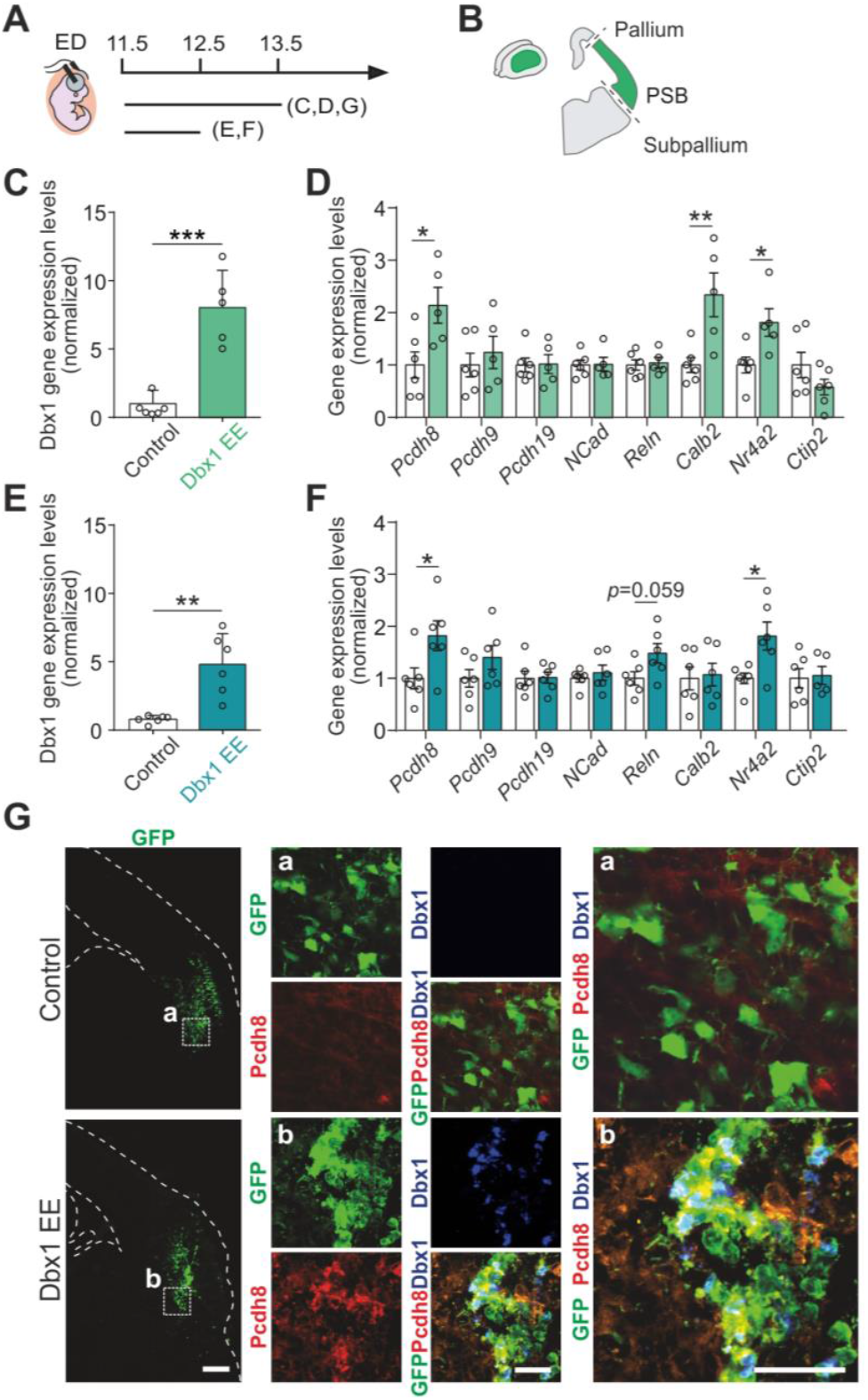
Dbx1 EE induces Pcdh8 gene expression in the electroporated area **(A)** Schematic timeline of the IUE. ED, electroporation day. **(B)** Electroporated region dissection scheme. **(C-F)** qPCR quantifications of **(C, E)** *Dbx1* or **(D, F)** *Pcdh8, Pcdh9, Pcdh19, Ncad, Reln, Calb2, Nr4a2* and *Ctip2* gene expression upon IUE at E11.5 of a control vector or Dbx1 EE and analysis at **(C, D)** E13.5 and **(E, F)** E12.5. Data are mean ± SEM; circles represent values from independent electroporations (*n*=6 controls, *n*=5-6 Dbx1 EE). Student’s *t*-test: **(C)** \*\*\**p*=0.0002, **(E)** \*\**p*=0.0014, **(D, F)** \**p*≤0.05 and \*\**p*≤0.01. **(G)** Confocal images of GFP (green) in coronal sections of E13.5 mouse brains electroporated at E11.5 with a control vector or Dbx1 EE, co-labeled with Pcdh8 (red) and Dbx1 (blue). Dashed squares (a, b) magnified on the right. Dashed lines outline the pallium and ventricle. Scale bars: 200 μm, 100 μm (magnified). See also Figures S5 and S6.

To validate these findings, and determine whether changes in gene expression are CA, we performed co-immunostaining of Pcdh8 and Dbx1 on E11.5 Dbx1-electroporated brain sections. We confirmed higher CA Pcdh8 protein expression in Dbx1 EE cells at E13.5 (Figure 3G). At E18.5, the picture was more complex as we observed both clusters of Pcdh8^high^Ctip2^−^Nr4a2^+^ cells below the CP, and Pcdh8^low^Ctip2^+^Nr4a2^−^ cell streams within the CP (Figure S6A and Figure 1I). Furthermore, following IUE at E11.5, the regular lateral Pcdh19 expression on E13.5 control sections was strongly reduced specifically where Dbx1-induced aggregates were formed (Figure S6B), consistent with the negative correlation with *Dbx1* gene expression (Figure S5A), supporting a *Dbx1*-mediated repression of *Pcdh19* transcription. Overall, these findings indicate a rapid and possibly time-restricted Dbx1 EE action on Pcdh8 upregulation as well as Pcdh19 repression in a CA manner.

Since aggregation and cell fate allocation differed between DP and LP/VP, and δ-Pcdh (i.e. Pcdh8, Pcdh9 and Pcdh19) exclusively mediate homophilic aggregation while avoiding heterotypic aggregation (Bisogni et al., 2018), we performed precise comparison of expression patterns by ISH for *Pcdh8, Pcdh9* and *Pcdh19* at E11.5, E12.5 and E13.5. We found that expression levels of different *Pcdh8, Pcdh9* and *Pcdh19* is distinct in LP *versus* DP. For example, *Pcdh19* is strongly expressed in progenitors, following a medial-to-lateral gradient with almost no expression at E12.5-E13.5, persisting only in postmitotic neurons of the future piriform cortex, whereas *Pcdh9* exhibits strong expression in the postmitotic compartment with a lateral-to-medial gradient and is highly expressed in progenitors around the PSB (Figure S5G-I). Moreover, the expression patterns for all three Pcdhs were extremely dynamic between E11.5 (IUE timepoint) and E13.5 (analysis timepoint). Thus, high levels of Pcdh8 in Dbx1 EE-expressing cells promoted strong homophilic interactions *in vivo* (aggregate formation) that occur preferentially in LP progenitor cells with low levels of other δ-Pcdh (i.e. Pcdh19). We conclude that dynamic developmental changes in adhesion molecule combinatorial code, progenitor *vs*. postmitotic identity and regional differences (medial-dorsal-lateral axis) (Figure S5G-I), modulate cell-segregative adhesion strength in different DV regions (Bisogni et al., 2018). These results thus reflect the important spatio-temporal changes that occur in the ability of the cortical neuroepithelium to respond to Dbx1 between E11.5 and E12.5 and point to possible distinct Pcdh and Cdh programs induced by Dbx1 at these stages.

### Pcdh8 is required for Dbx1-induced adhesion and fate specification

To determine the contribution of Pcdh8 to Dbx1-induced cell aggregation, we further investigated the epistasis between Dbx1 and Pcdh8 by combining Dbx1 EE with *Pcdh8* knock-down (KD). We first designed three shRNA constructs targeting *Pcdh8* and determined their efficiency by western blot analysis 48h after co-transfection with a *Pcdh8* expression vector in HEK293T cells. We selected the most effective shRNA, which reduced *Pcdh8* expression by 51% compared to a control shRNA (Figure S7A, C). When electroporated alone, the Pcdh8 shRNA did not induce obvious changes in cell aggregation (Figure S7D) or cell identity (Nr4a2, Ctip2) (Figure S7E, F). Next, we co-electroporated E11.5 mouse brains with Dbx1 EE and Pcdh8 shRNA or control shRNA constructs (Figure 4A, B, and C). Interestingly, we observed no aggregate formation with Pcdh8 KD (Figure 4B, C), thus indicating that Pcdh8 is required for Dbx1-induced adhesion. Moreover, we observed a reduced number of GFP^+^Nr4a2^+^ cells (Figure 4D), and an increased number of GFP^+^Ctip2^+^ cells (Figure 4E) compared to Dbx1 EE + control shRNA or Dbx1 EE alone, indicating that Pcdh8 is also necessary for Dbx1-induced fate determination. Since Pcdh8 KD rescued the phenotypes induced by Dbx1 EE, we decided to monitor Dbx1 expression upon Pcdh8 KD. Surprisingly, we observed numerous GFP^+^Dbx1^−^ cells (Figure 4C, F, G) in comparison to Dbx1 EE alone (Figure 3G, Figure S1C, D). Since Dbx1 EE is driven by a plasmid bearing a heterologous CAG promoter, Pcdh8 shRNA likely affects Dbx1 expression post-transcriptionally.

**Figure 4.**
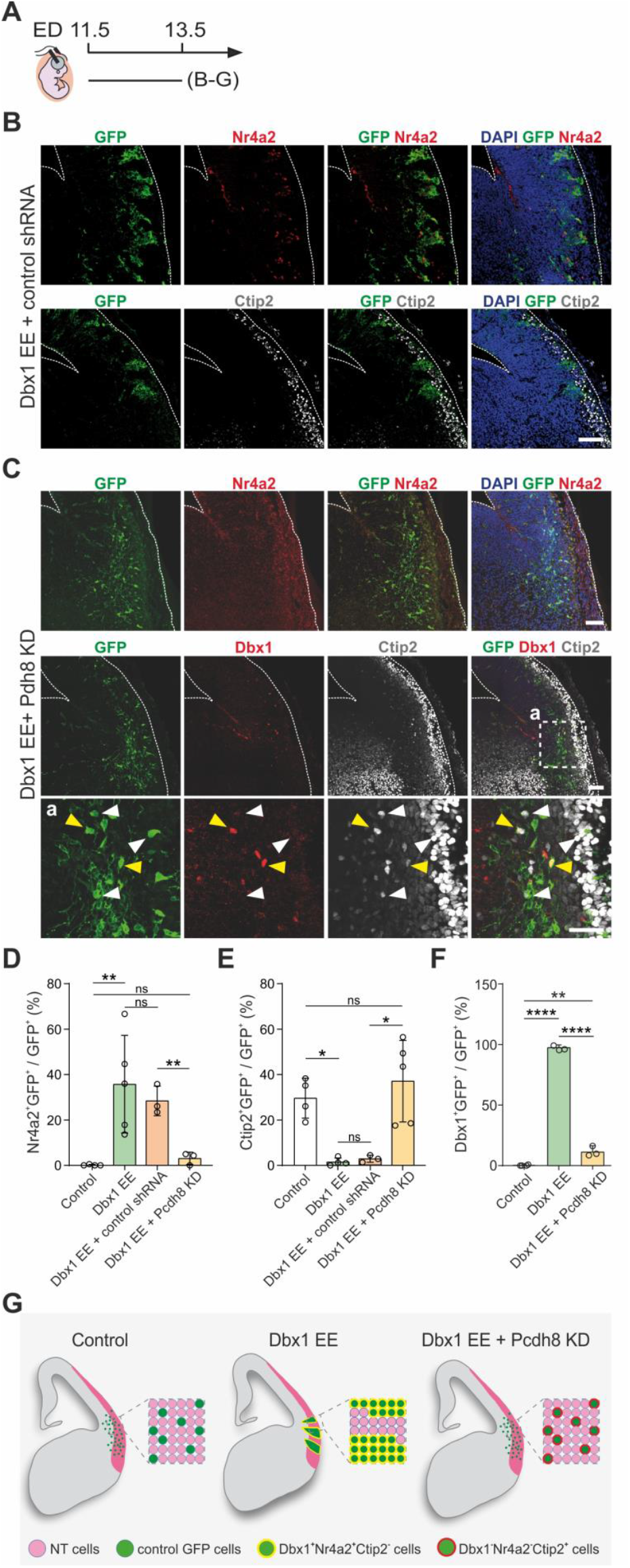
Pcdh8 is responsible for the Dbx1 EE phenotype. **(A)** Schematic timeline of the IUE. ED, electroporation day. **(B, C)** Confocal images of GFP (green) in coronal sections of E13.5 mouse brains electroporated at E11.5 with Dbx1 EE + control shRNA or Dbx1 EE + Pcdh8 KD (Pcdh8 shRNA), co-labeled with **(B, C)** Nr4a2 (red) or **(B)** Ctip2 (gray), or with **(C)** Dbx1 (red) and Ctip2. Dashed square (a) magnified below. White and yellow arrowheads point to Dbx1^−^ and Dbx1^+^ cells, respectively. Dashed lines outline the pallium and ventricle. Scale bars 100 μm. **(D-F)** Percentages of **(D)** Nr4a2^+^GFP^+^, **(E)** Ctip2^+^GFP^+^ and **(F)** Dbx1^+^GFP^+^ cells among GFP^+^ cells. Data are mean ± SEM; circles represent values from independent electroporations (*n*=4 controls, *n*=3-5 Dbx1EE, *n*=3 Dbx1 EE + control shRNA, *n*=3-5 Dbx1 EE + Pcdh8 KD). One-way ANOVA, *post hoc* Holm-Sidak: ns, not significant, **(D)** \*\**p*=0.0099 and \*\**p*=0.0034, **(E)** \**p*<0.05 \**p*=0.0187 **(F)** \*\*\*\**p*<0.0001 and \*\**p*=0.0021. **(G)** Scheme of Pcdh8 KD effect on Dbx1 EE-induced aggregate formation phenotype and cell fate. See also Figure S7.

We thus conclude that Pcdh8 is an essential component in Dbx1-induced aggregation and cell-fate determination.

### Pcdh8 EE results in major reorganization of pallial domains

To further investigate the epistasis between Dbx1 and Pcdh8, we performed Pcdh8 gain-of-function (Pcdh8 EE) in the LP, which is a territory devoid of Dbx1 expression, by IUE at E11.5 (E11.5-13.5) using a pCAGGS-Pcdh8-iresGFP plasmid. First, we confirmed Pcdh8 ectopic expression through western blotting (Figure S7A, B) and immunofluorescence (Figure S7G). In addition, ISH revealed similar expression levels of *Dbx1* and *Pcdh8* in both Dbx1 EE and Pcdh8 EE experiments (Figure S7H). Upon Pcdh8 EE, we observed that the Pcdh8 resulted in cortical plate thickening compared to the contralateral non-electroporated hemisphere (Figure 5A-C). ISH experiments revealed major tissue reorganization, consisting in the formation of large circumferential GFP^+^ structures that recapitulated the DV partitioning of pallial domains with patches or stripes of cells expressing the ventral markers *Shh* or *Gsx2* segregated from cells positive for the dorsal markers *Lhx2* or *Bhlhe22* (Figure 5D and Figure S7I, J). Moreover, we observed that, markers of cycling progenitors such as Pax6 (Figure 5E, F) or *Notch1* (Figure 5D and Figure S7J) were affected and extended outside of the VZ. Strikingly, *Dbx1*, which is normally restricted mostly to BPs of the VP, was found expressed throughout the electroporated area (Figure 5D and Figure S7J). Altogether, Pcdh8 EE caused the local overgrowth of rosette-like structures, characterized by an outer layer of Pcdh8, *Dbx1, Bhlhe22* and *Shh*-expressing cells, segregating and organizing *Pax6, Lhx2* and *Notch1*-positive cells at the centre. The rosettes often displayed a central lumen lined by PH3^+^ mitotic cells, surrounded by GFP^+^ (Pcdh8^+^) cells (Figure 5G, H) and delineated by Ncad^+^ apical junctions (Figure 5I). We observed the delamination of PH3^+^ APs (Figure 5G) that correlated with the disruption of Ncad^+^ adherens junctions in the neuroepithelial cells lining the ventricle (Figure 5I).

**Figure 5.**
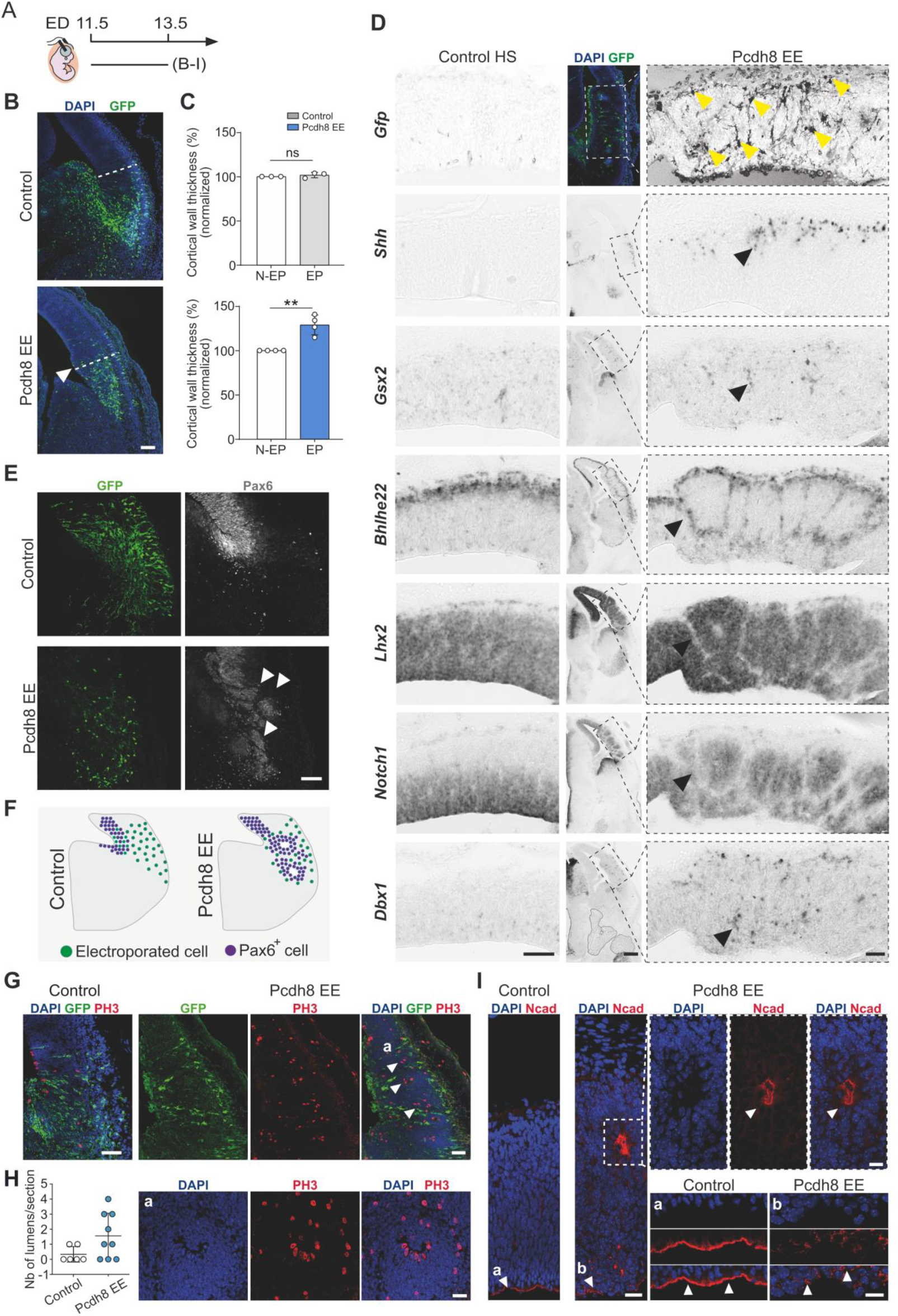
Pcdh8 EE causes reorganization of the developing pallium. **(A)** Schematic timeline of the IUE. ED, electroporation day. **(B)** Confocal images of GFP (green) in coronal sections of E13.5 mouse brains electroporated at E11.5 with control or Pcdh8 EE, with DAPI (blue) counterstaining. White arrowhead indicates the region of developing pallium overgrowth. Dashed lines show the location of thickness measurements. Scale bar: 200 μm. **(C)** Quantification of cortical wall thickness from **(B)** in electroporated (EP) control vector (top) or Pcdh8 EE (bottom) normalized to the thickness of the non-electroporated (N-EP) contralateral hemisphere. Data are mean ± SEM; circles represent values from independent electroporations (*n*=3 controls, *n*=4 Pcdh8 EE). Student’s *t*-test: \*\**p*=0.0020, ns, not significant. **(D)** Confocal image of GFP (green) in the Pcdh8 EE electroporated area (top middle panel), counterstained with DAPI (blue), and bright-field images of *in situ* hybridization for *Gfp, Shh, Gsx2, Bhlhe22, Lhx2, Notch1* and *Dbx1* in coronal sections of E13.5 mouse brains electroporated at E11.5. The left panel shows the control contralateral hemisphere (HS) from the same section as Pcdh8 EE (right panel). Dashed rectangles from the Pcdh8 EE area magnified on the right. Yellow arrowheads indicate cells displaying Pcdh8 EE. Black arrowheads indicate altered mRNA expression. Scale bars: 200 μm, 100 μm (magnified). **(E)** Confocal images of GFP (green) in coronal sections of E13.5 mouse brains electroporated at E11.5 with control vector or Pcdh8 EE, co-labeled with Pax6 (gray). White arrowheads indicate the disorganization of Pax6^+^ cells within the ventricular zone (VZ) and ventral pallium (VP). Scale bar: 100 μm. **(F)** Scheme of Pax6^+^ cells distribution 48h after IUE as in **(E). (G)** Confocal images of GFP (green) in coronal sections of E13.5 mouse brains electroporated at E11.5 with control vector or Pcdh8 EE, co-labeled with PH3 (red) and DAPI (blue) counterstaining. White arrowheads indicate PH3^+^ cells outside the VZ. Scale bar: 100 μm. Rosette (a) magnified at the bottom. Scale bar: 50 μm. **(H)** Quantification of rosettes in control and Pcdh8 EE brain sections. Data are mean ± SEM; circles represent values of single sections (*n*=6 controls, *n*=9 Pcdh8 EE). **(I)** Confocal images of representative regions from control and Pcdh8 EE (IUE E11.5-E13.5) labeled with Ncad (red) and DAPI (blue) counterstaining. Dashed square and indicated VZ (a, b) Magnified at the top right and bottom right, respectively. White arrowheads indicate disturbance of Ncad expression. Scale bars: 100 μm (left panel), 50 μm (right panels). See also Figure S7.

We therefore conclude that ectopic Pcdh8 expression disrupts the local organization of the telencephalic neuroepithelium through the modulation of cell adhesion, resulting in alterations in DV patterning gene expression and apico-basal polarity within the LP.

### Cell-autonomous control of cell fate by Pcdh8

The very precise segregation of gene expression in outer and central domains of the observed rosette-like structures prompted us to evaluate more precisely the CA *versus* NCA function of Pcdh8. We initially performed Dbx1 IF on brain sections of E11.5-electroporated embryos, 48h after IUE with Pcdh8 EE (E11.5-E13.5). The analysis revealed almost complete co-localization, with 90% ± 5% of Pcdh8 EE-expressing cells (GFP^+^) being Dbx1^+^ compared to 0% of control GFP-transfected cells (Figure 6A-C). Furthermore, we observed that 16.47% ± 2.38% of Pcdh8 EE-expressing cells were Nr4a2^+^Ctip2^−^, compared to only 0.29% in controls (Figure 6D, E), while the Nr4a2^+^Ctip2^+^ SP-like cells remained low and comparable between Pcdh8 EE and control conditions (Figure 6D, F). This suggests that, similar to Dbx1 EE, Pcdh8 EE does not produce SP neurons in the LP. Additionally, the proportion of Ctip2^+^Nr4a2^−^ cells was reduced among Pcdh8-transfected cells (19.92 ± 0.95%) compared to controls (29.54 ± 4.37%) (Figure S8A, B). These findings are consistent with the cell fate effects of Dbx1 EE (Figure 1), indicating that Pcdh8 can regulate a similar cell fate, likely of the CLA lineage (Nr4a2^+^Ctip2^−^), in the developing LP in a CA manner.

**Figure 6.**
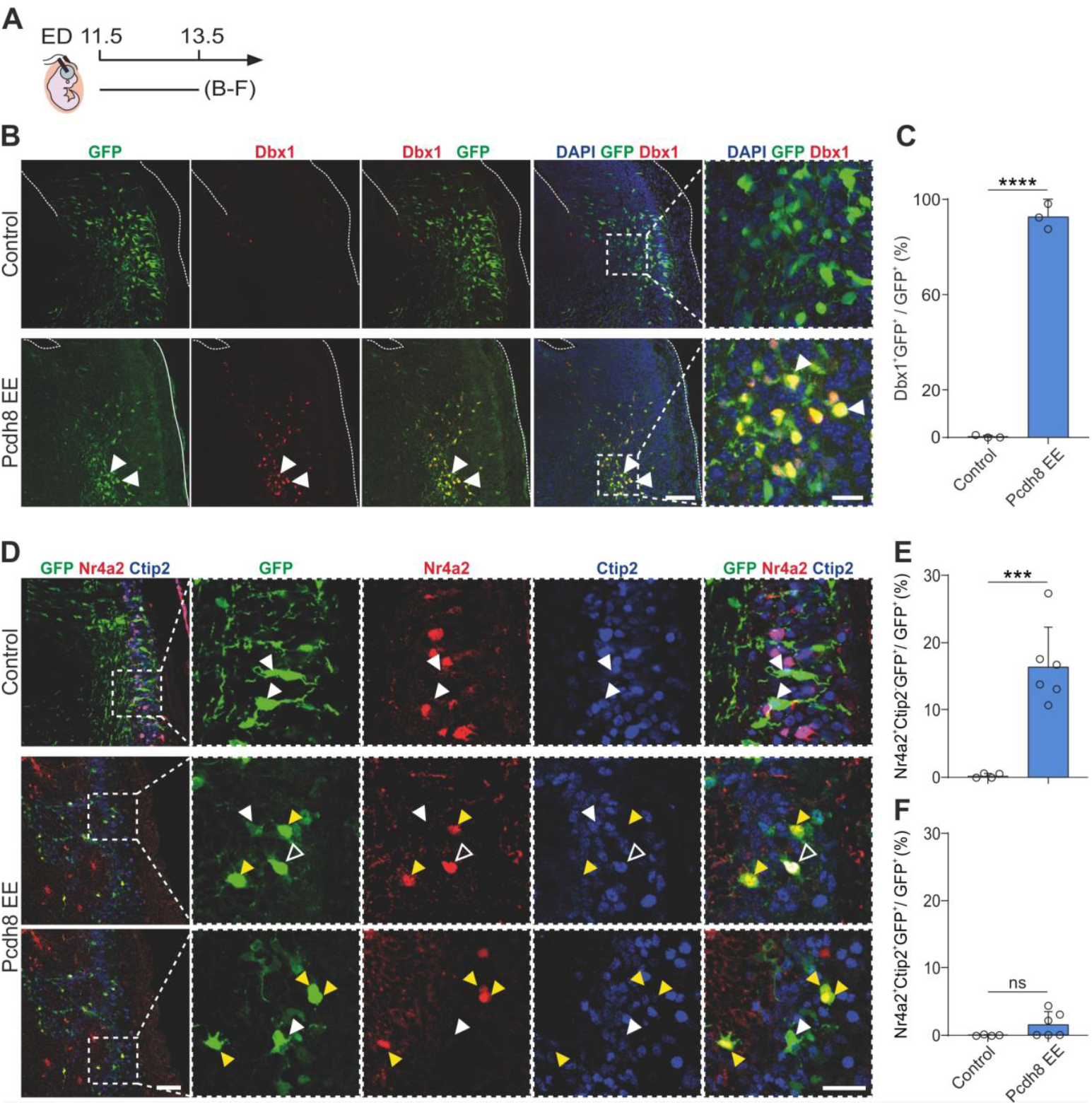
Pcdh8 EE induces Dbx1 and Nr4a2^+^ CLA neuron fate. **(A)** Schematic timeline of the IUE. ED, electroporation day. **(B)** Confocal images of GFP (green) in coronal sections of E13.5 mouse brains electroporated at E11.5 with control vector or Pcdh8 EE, co-labeled with Dbx1 (red) and DAPI (blue) counterstaining. White arrowheads indicate Dbx1^+^GFP^+^ cells. Dashed squares highlighting GFP^+^ regions magnififed on the right. Dashed lines outline the pallium and ventricle. Scale bars: 200 μm, 50 μm (magnified). **(C)** Percentage of Dbx1^+^GFP^+^ cells over the total of GFP^+^ cells. Data are mean ± SEM; circles represent values from independent electroporations (*n*=3 controls, *n*=3 Pcdh8 EE). Student’s *t*-test: \*\*\*\**p*< 0.0001. **(D)** Confocal images of GFP (green) in coronal sections of E13.5 mouse brains electroporated at E11.5 with control vector or Pcdh8 EE, co-labeled with Nr4a2 (red) and Ctip2 (blue). Dashed boxes magnified on the right. Yellow, empty and filled white arrowheads indicate Nr4a2^+^Ctip2^−^GFP^+^, Nr4a2^+^Ctip2^+^GFP^+^ and Ctip2^+^Nr4a2^−^GFP^+^ cells, respectively. Scale bars: 100 μm, 50 μm (magnified). **(E, F)** Percentages of **(E)** Nr4a2^+^Ctip2^−^ GFP^+^ and **(F)** Nr4a2^+^Ctip2^+^GFP^+^ cells among GFP^+^ cells. Data are mean ± SEM; circles represent values from independent sections (*n*=3 electroporations/condition, 1-2 sections/electroporation). Student’s *t*-test: **(E)** \*\*\**p*=0.0006; **(F)** ns, not significant. See also Figure S8.

To further investigate the mechanisms by which Pcdh8 controls the establishment of neuronal identity, we monitor cell proliferation and cell cycle exit following Pcdh8 IUE. We first quantified the number of mitotic progenitors among transfected cells (GFP^+^PH3^+^) and compared their positioning within the neuroepithelium of Pcdh8- and control GFP-electroporated brains. We observed a complete CA switch from ventricular to abventricular mitotic figures upon Pcdh8 EE (Figure S8C-F), consistent with the observed disruption of apico-basal polarity (Figure 5D, G, I). Additionally, after the administration of a single pulse of EdU at E12.5, thus 24h post-surgery, quantification of EdU^+^ transfected cells revealed that Pcdh8 EE led to a ~50% reduction in S-phase entry compared to controls (Figure S8A, C, G) suggesting premature cell cycle exit.

Given the massive disruption of the mitotic/postmitotic compartmentalization upon Pcdh8 EE (Figure 5D-H and Figure S7J) and the role of the Notch pathway in maintaining the radial glia phenotype (Blackwood, 2019) and preventing premature differentiation (Ramos et al., 2010), we examined the expression of Notch ligands Delta1 and Jag1 (Lasky and Wu, 2005; Stump et al., 2002) 48h following Pcdh8 IUE of E11.5 mouse brains (E11.5-E13.5) (Figure 7A). We found that a NCA Delta1 upregulation at the centre of Pcdh8-induced rosettes (Figure 7B) is accompanied by a CA decrease in Jag1 expression (Figure 7C). To functionally assess whether Jag1 downregulation is responsible for the observed phenotype, we performed co-electroporation of Pcdh8 EE with a Jag1-expressing plasmid. We showed that Jag1 EE prevented the formation of rosettes, as well as the associated overgrowth and abnormal cell distribution, effectively rescuing the phenotype (Figure 7D). Together, the differential regulation of Delta1 and Jag1 by Pcdh8 EE, along with the rescue of the phenotype by Jag1 EE, indicate that the Notch pathway mediates the response to Pcdh8. These findings suggest that Jag1 downregulation is responsible for altering the fate program of transfected progenitors (Figure 7E) and disrupting the neuroepithelial organization (Figure 7F).

**Figure 7.**
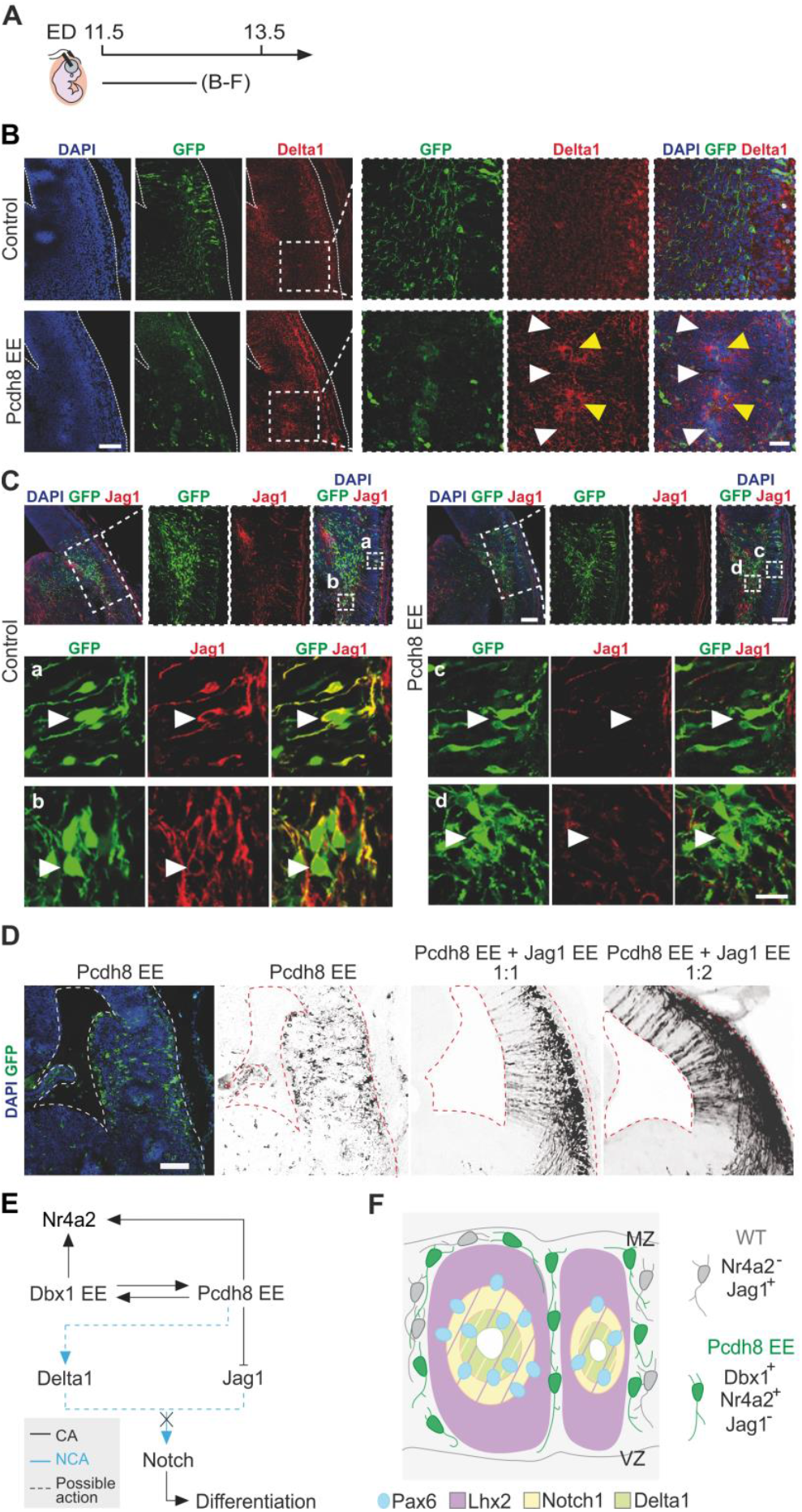
Pcdh8 EE phenotype is mediated by the Notch1 pathway. **(A)** Schematic timeline of the IUE. ED, electroporation day. **(B)** Confocal images of GFP (green) in coronal sections of E13.5 mouse brains electroporated at E11.5 with control vector or Pcdh8 EE, co-labeled with Delta1 (red) and DAPI (blue) counterstaining. Scale bar: 200 μm. Dashed squares magnified on the right. Yellow and white arrowheads indicate Delta1 accumulation signal or surrounding GFP^+^ cells with low Delta 1 signal, respectively. Dashed lines outline the pallium and ventricle. Scale bar: 50 μm. **(C)** Confocal images of GFP (green) in coronal sections of E13.5 mouse brains electroporated at E11.5 with control vector or Pcdh8 EE, co-labeled with Jag1 (red) and DAPI (blue) counterstaining. Dashed rectangles magnified on the right. Scale bars: 200 μm, 100 μm (magnified). Dashed squares in control (a, b) and Pcdh8 EE (c, d) magnified in the bottom. White arrowheads indicate representative cells of each condition. Scale bar: 25 μm. **(D)** Confocal images of coronal sections of E13.5 mouse brain cortices electroporated at E11.5 with either Pcdh8 EE alone or co-electroporated with Jag1 EE GFP (green) and DAPI (blue) staining are shown, as well as black-and-white images of GFP expression for better visualization. Dashed lines delineate the pallium and ventricle. Scale bar: 100 μm. **(E)** Scheme of the possible interactions between Dbx1, Pcdh8, Notch signaling and cell fate markers. CA, cell autonomous; NCA, non-cell autonomous. **(F)** Cartoon showing composition of the rosette-like structures formed upon Pcdh8 EE.

Taken together, these results show that manipulating cell adhesion through ectopic Pcdh8 expression is sufficient to affect neuronal fate acquisition in a CA manner *via* Notch signalling alteration and premature cell cycle exit.

## DISCUSSION

### Adhesion is an important component of cell identity

TFs accomplish a key role in the formation and maintenance of different cell types during development. Indeed, changes in their expression levels (e.g. due to ectopic expression of Pax6, Lhx2 or Dbx1) alter initial programs of progenitor cells (Arai et al., 2019; Pataskar et al., 2016; Subramanian et al., 2011). Our data demonstrate that changes in Dbx1 expression influence cellular adhesive properties *via* Pcdh8, making it an essential element of cell identity that is sensitive to Pcdh levels. Interestingly, an emerging role of cPcdhs in spatial and functional organization of neurons has recently been shown in the neocortex (Lv et al., 2022). Indeed, it has been demonstrated that cPcdhs are expressed in patterned combinations by excitatory neurons that arise from the same progenitor cell. Furthermore, a specific surface pattern of cPcdhs is important for neuron positioning and connectivity with other neurons (Lv et al., 2022). Nevertheless, the regulatory mechanism regulating the expression of distinct cPcdhs gene patterns has not been described. Here, we demonstrate that EE of the TF Dbx1 changes the spatiotemporal competence of the cortical neuroepithelium. These observations are consistent with the established role of endogenous Dbx1 as a strong cell-fate determinant in different CNS regions (Arai et al., 2019; Bouvier et al., 2010b; Pierani et al., 2001; Sokolowski et al., 2016). Specifically, we found that Dbx1 EE imparts different cell fates in a time- and region-dependent manner. Temporally, it promoted the generation of Nr4a2^+^Ctip2^−^ cells (not SP neurons) at E11.5 and Nr4a2^−^ Ctip2^+^ at E12.5. Regionally, at E11.5, it gives rise to CRs and SP neurons in the DP (Arai et al., 2019), while in the LP/VP it induced Nr4a2^+^Ctip2^−^ cells characteristic of the CLA (Puelles et al., 2016b; Bruguier et al., 2020; Mantas et al., 2024; Yan et al., 2025). This correlates with the induction of distinct set of Pcdhs, namely Pcdh8 at E11.5 and Pcdh9/Pcdh19 at E12.5, suggesting that progenitor cell competence varies over time, giving rise to daughter cells with different Pcdhs expression combinations on their surface. In our study, Dbx1 EE in the LP led to the organization of transfected cells into Pcdh8-expressing aggregates with CLA neuron characteristics (Nr4a2^+^Ctip2^−^Reln^−^Calb2^−^), whereas in the DP, it did not induce aggregation and instead correlated with CR and SP neuron fates (Arai et al., 2019). We believe that the strong cell-segregative character of Pcdh8 and the quick changes in Pcdhs expression patterns on the DV axis can explain the differences in aggregate formation between the dorsal and lateral pallium. Variations in adhesion properties can play a crucial role in segregating and organising cell types into properly functioning brain areas. Moreover, Pcdh8 EE resulted in the reorganization of multiple TF expression crucial for correct DV patterning (e.g. *Shh, Dbx1, Lhx2*). Notably, during early stages of spinal cord development, specific cell adhesion molecules have been shown to follow the expression of TFs in selected progenitor domains (Tsai et al., 2020), ensuring robust tissue patterning. Altogether, our findings suggest that Dbx1 expression is both temporally and spatially restricted, and together with Pcdh8, can influence TF spatial organization, thereby contributing to proper cortical development.

### Crosstalk between Pcdh8 and TFs determines CLA fate

Dbx1 was previously shown to function as a potent determinant of cell identity in the mouse spinal cord (Pierani et al., 2001). During early development, Dbx1-expressing domains are present at the borders of the pallium and subpallium, notably the VP/PSB and septum (Bielle et al., 2005; Griveau et al., 2010). Interestingly, we detected co-expression of Dbx1 and Pcdh8 in VP and septal progenitors, and Dbx1-derived cells in the postmitotic compartment of the septum and VP in a ventrally migrating stream toward the prospective CLA. We showed changes in *Pcdh8* expression domains in Dbx1^*LacZ/LacZ*^ KO mice, suggesting that Dbx1 regulates Pcdh8 expression in a region-specific manner, restricting its expression in septal progenitors while promoting its expression in the VP. This pattern correlates with the emergence of the lateral postmitotic Pcdh8^+^ (Nr4a2^+^) population. Moreover, genetic lineage tracing in the developing mouse cortex demonstrated that Dbx1 progenitors also give rise to Nr4a2^+^ and Pcdh8^+^ neurons located in territories corresponding to the future CLA. These data argue in favour of a potential direct, though not exclusive, interaction between Dbx1 and Pcdh8 in cell identity determination. At the molecular level, Pcdh8 KD in Dbx1 EE cells prevented Dbx1 expression and consequently reversed CLA cell identity, whereas Pcdh8 EE induced both Dbx1 expression and CLA fate. While Dbx1 may control Pcdh8 expression at the transcriptional level, we propose that Pcdh8 act as a post-transcriptional modulator of Dbx1, since the same pCAGGS promoter drives the expression in both EE and KD experiments. Together, these results clearly show that Dbx1 and Pcdh8 interact in a reciprocal manner within the VP/LP to specify CLA neuron identity. Complementing the findings of Arai et al. (2019), our data confirm that in mice, Dbx1^+^ progenitors do not generate SP neurons, unlike in primates, and identify the bidirectional molecular interaction between Dbx1 and Pcdh8 that mediates CLA-Nr4a2^+^ neuronal fate. This provides a previously unrecognized and important role of adhesion molecules in regulating TF expression and cell fate. Given the critical functions of the CLA region, including Nr4a2^+^ neurons, in attention control, slow wave sleep, depressive-like behavior, and pain processing (Puelles et al., 2016b; Bruguier et al., 2020; Mantas et al., 2024; Yan et al., 2025), our findings provide key insights into cortical development and functional organization.

### Pcdh8 regulate cortical development from cell to tissue organization levels

Multiple studies have shown that disturbance of TF expression, i.e. Lhx2 or/and Pax6, results in re-patterning of TF expression domains (Mangale et al., 2008; Roy et al., 2013; Godbole et al., 2017) and tissue overgrowth (Mangale et al., 2008). We found that Dbx1 EE influenced cell fate by inducing Pcdh8 expression without major disturbance of DV patterning. However, shifting Pcdh8 expression, typically restricted to the septum and/or ventrolateral domains, into more dorsal pallial regions disrupted the expression of TFs *Lhx2* and Pax6 in progenitor cells. This was accompanied by AP delamination due to the loss of Ncad^+^ adherens junctions in the neuroepithelial cell lining the ventricle and the formation of rosette-like structures containing new proliferative centres enriched with Ncad. Moreover, we observed ectopic expression of the subpallial progenitor TF marker *Gsx2* into the DP, while the pallial postmitotic *Bhlhe22* localized to the periphery of the rosettes, both in a CA or NCA manner. These observations are more profound but still consistent with studies on disruptions of the Gsx2-Pax6-Dbx1 border at the PSB (Cocas et al., 2011) or the hem-antihem balance due to Lhx2 dysregulation, which resulted in dramatic impairments in DV organization and, consequently, deformation of the developing pallium (Godbole et al., 2017). Furthermore, since Pcdh expression is confined to the apical membrane (Lobas et al., 2012) and Ncad is crucial for neuroepithelial integrity and cortical organization (Kodawaki et al., 2007), our work adds Pcdh8 as a new organizer of the DV axis. The observed differences likely derive from the timing of Pcdh8 expression, whether directly driven by the pCAGGS promoter/enhancer or induced later by Dbx1 EE (itself being first expressed under pCAGGS control). As Dbx1 promotes cell cycle exit (García-Moreno et al., 2018), pCAGGS-driven Pcdh8 will induce expression directly in APs, whereas pCAGGS-driven Dbx1 expression in APs will trigger rapid cell cycle exit, thus likely inducing Pcdh8 expression in young neurons (or BPs). Mechanistically, Pcdh8 might act on TF expression through the activation of intracellular signalling cascades *via* its intracellular domain (ICD). Specifically, the *cis*-binding of Pcdh8 to Ncad, where the Pcdh8 ICD activates the MAP kinase (MAPK) TAO2β, has been previously described (Yasuda et al., 2007). Interestingly, MAPK are key mediators of eukaryotic transcriptional responses to extracellular stimuli (Zarubin and Han, 2005; Whitmarsh, 2007). Additionally, Pcdh8 may exert post-transcriptional effects, but its specific impact in modulating Dbx1 expression remains unclear. Current literature suggests that this effect could occur through interactions with the TAO2β–p38-MAPK (Gao et al., 2022; Mahtani et al., 2001; Yasuda et al., 2007) or GSK3/β-catenin pathways (Farina et al., 2009; Gizak et al., 2020; Unterseher et al., 2004; Zong et al., 2017), both of which regulate gene expression at the post-transcriptional level. Therefore, it would be interesting to further investigate the action of Pcdh8 in both transcriptional and post-transcriptional mechanisms. Collectively, these results demonstrate that Pcdh8, which is normally strongly expressed in progenitors at the septum and VP/PSB, when ectopically expressed at or dorsal to the PSB can significantly affect DV organisation and the activity of several TFs in both CA and NCA manners.

### Notch signalling as a downstream partner of Pcdh8

The Notch signalling pathway tightly controls both the temporal and spatial patterning of neuronal populations (Wang et al., 2015; Ware et al., 2016). Inactivation of Notch1 has been associated to accelerated neuronal differentiation in the mouse spinal cord and developing brain (Kong et al., 2015; Tokunaga et al., 2004; Yang et al., 2006). Additionally, it controls the number of neural progenitor cells that exit the cell cycle and undergo differentiation (Moore and Alexandre, 2020). We showed that Pcdh8 influences cell cycle exit and postmitotic fate acquisition. Moreover, we found that Pcdh8 represses Notch1 signalling ligands Jag1 and Dll1 in CA and NCA manners, respectively, thus suggesting a potential impact on differentiation. A rescue experiment combining Pcdh8 EE with Jag1 EE further demonstrated that restoring Jag1 in Pcdh8 EE restored normal pallial formation, providing strong evidence that Jag1 downregulation is responsible for the Pcdh8 EE-induced phenotype. Interestingly, Dbx1 was also shown to influence Jag1 expression in the spinal cord (Skaggs et al., 2011) and cell cycle exit (García-Moreno et al., 2018). Together, these findings position the Notch pathway as a relevant candidate for the cooperative function of Pcdh8 and Dbx1, which pushes progenitors out of the cell cycle toward premature differentiation.

In summary, our study establishes adhesion molecules, notably Pcdh8, as novel players in pallium morphogenesis and cell fate determination. Above all, we demonstrate that Pcdh8, through a bidirectional regulatory mechanism with the TF Dbx1, governs cell cycle dynamics and fate acquisition, ultimately shaping the organization of the developing brain. Moreover, by identifying Notch signaling as a downstream effector of Pcdh8, our findings sheds light on the regulatory mechanism behind these processes, providing new insights into the molecular orchestration of neural development.

## RESOURCE AVAILABILITY

### Lead contact

Further information and requests for resources or raw data should be directed to the lead contacts, Alessandra Pierani and Andrzej W. Cwetsch.

### Materials availability

Shiny App to explore scRNAseq data from the mouse septum is available at: https://apps.institutimagine.org/mouse_septum/. Additionally, the Shiny App that allows making the pseudotime reconstructions of the ventral pallium (Moreau et al., 2021) is available at: https://apps.institutimagine.org/mouse_pallium/.

### Data and code availability

Raw scRNAseq reads and processed count matrix are available from GEO (accession number GSE229603). The barcodes, SPRING coordinates and metadata of cells retained after quality control, as well as annotated R codes have been deposited at https://fcauseret.github.io/septum/.

## ACKNOWLEDGMENTS

We thank E. Panafieu, Animalliance and the Imagine Institute animal house staff for mouse care and technical help. We also appreciate advice in cloning process and donation of pStrike plasmid by P. Billuart. Furthermore, we thank Stéphane Nedelec, Sophie Thomas, Jose Manuel Morante Redolat and Isabel Martinez Garay for their help in the experiments not included in this manuscript. Finally, we thank Isabel Fariñas for letting the laboratory space to perform some of the experiments and Cristina Gil-Sanz for Tle4 antibody. This work was supported by Fondation pour la Recherche Médicale; Number of the project: SPF20170938863, Fundation Fyssen; Number of the project: CH9147, Generalitat Valenciana; Number of the project: CDEIGENT/2021/005 to A.W.C., by grants from the Agence Nationale de la Recherche (ANR-2011-BSV4-023-01, ANR-15-CE16-0003-01 and ANR-20-CE16-0001-01), FRM (Equipe FRM DEQ20130326521 and EQU201903007836) to A.P; and state funding from the Agence Nationale de la Recherche under ‘‘Investissements d’avenir’’ program (ANR-10-IAHU-01) to the Imagine Institute A.P. is a CNRS (Centre National de la Recherche Scientifique) Investigator.

## CO-AUTHORS CONTRIBUTION

Conceptualization, A.W.C., F.C., A.P.; Methodology A.W.C., F.C. and A.P.; Investigation, A.W.C., S.F., E.D., J.G.J., M.X.M., Y.S., P.G.B., S.C.P., J.G.M., U.B.; Data Curation, A.W.C.; Writing – Original Draft, A.W.C. and A.P.; Writing – Review & Editing, A.W.C., S.F., F.C. and A.P.; Visualization, A.W.C., S.F., M.X.M. and F.C.; Supervision, A.P.; Project Administration, A.W.C. and A.P.; Funding Acquisition, A.W.C and A.P.

## SUPPLEMENTAL INFORMATION

**Figure S1.**
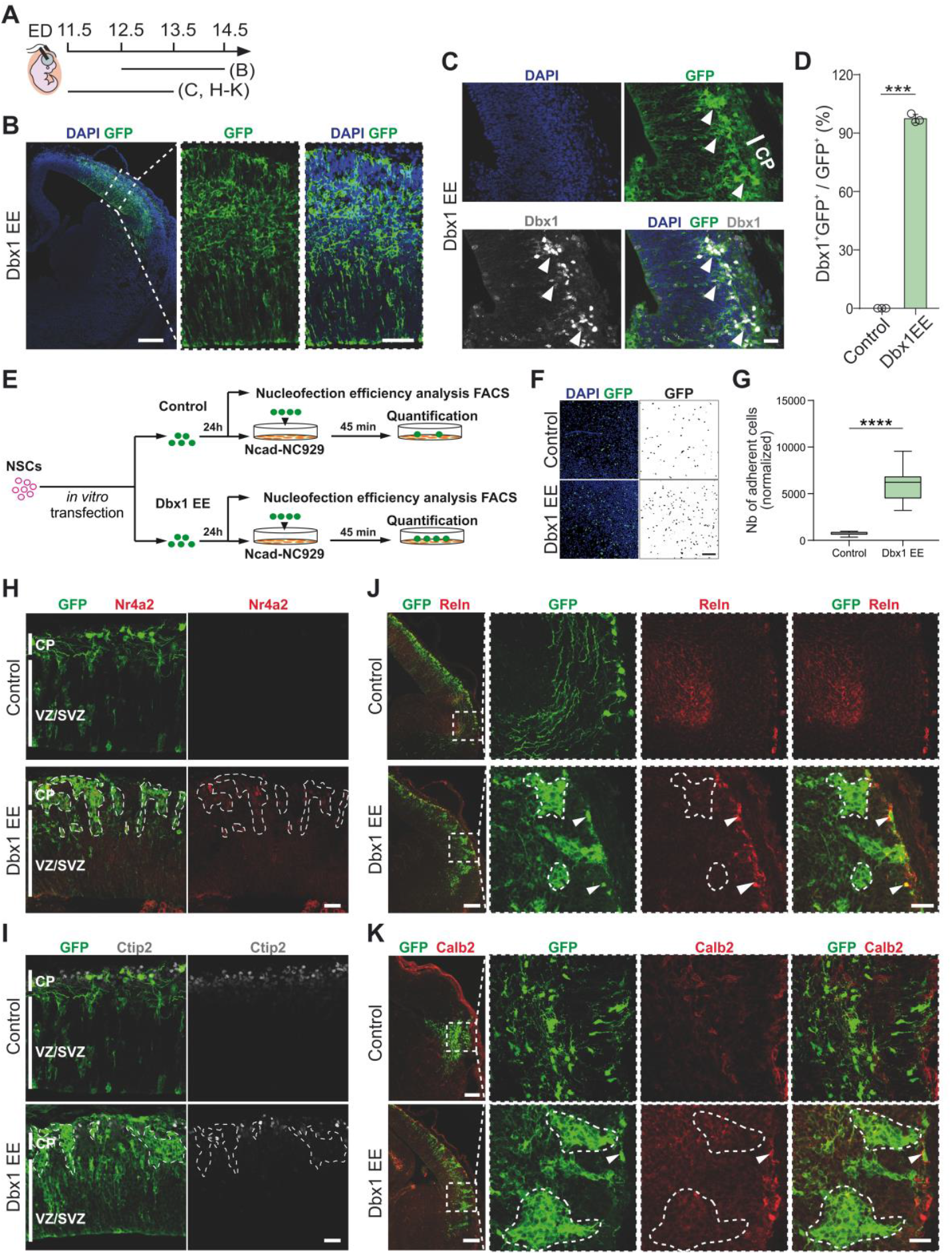
Dbx1 EE at E11.5 induces the formation of cell aggregates composed of SP-like neurons. (A) Schematic timeline of the IUE. ED, electroporation day. (B) Confocal images of GFP (green) in coronal sections of E14.5 mouse brains electroporated at E12.5 with Dbx1 EE, with DAPI (blue) counterstaining. Dashed rectangle magnified on the right. Scale bars: 200, 100 μm (magnified). (C) Confocal images of GFP (green) in coronal sections of E13.5 mouse brains electroporated at E11.5 with Dbx1 EE, co-labeled with Dbx1 (gray) and DAPI (blue) counterstaining. White line shows the cortical plate (CP). White arrowheads indicate Dbx1+GFP+ cells. Scale bar: 50 μm. (D) Percentage of Dbx1+GFP+ cells among GFP+ cells. Data are mean ± SEM; circles represent values from independent electroporations (n=3 each condition). Student’s t-test: ***p<0.001. (E) Schematic representation of the in vitro adhesion assay. Twenty-four hours after transfection, neural stem cells (NSCs) transfected with either a control vector or Dbx1 EE were analyzed by FACS to assess nucleofection efficiency, or seeded onto Ncad-expressing NC929 cell monolayers for 45 min before quantification. (F) Confocal images of transfected GFP+ (green) cells with a control vector or Dbx1 EE plasmid, with DAPI (blue) counterstaining. Black-and-white images of adherent GFP+ cells are show in the right panel. Scale bar: 100 μm. (G) Quantification of the number of adherent cells transfected with control or Dbx1 EE vector normalized to the percentage of all GFP+ detected by FACS in each corresponding condition. Student’s t-test: ****p≤0.0001. (H, I) Confocal images of GFP (green) in coronal sections of E13.5 mouse brains electroporated at E11.5 with control vector or Dbx1 EE, co-labeled with (H) Nr4a2 (red) and (I) Ctip2 (gray). White lines show the CP and the ventricular/subventricular zones (VZ/SVZ). Scale bars: 50 μm. (J, K) Left: Confocal images of GFP (green) in coronal sections of E13.5 mouse brains electroporated at E11.5 with control vector or Dbx1 EE, co-labeled with (J) Reln (red) or (K) Calb2 (red). Scale bars: 200 μm. Right: Dashed squares magnified. Scale bars: 50 μm. White arrowheads indicate (J) GFP+Reln+ and (K) GFP+Calb2+ cells localized outside the aggregates. Dashed areas delineate Dbx1 EE-induced aggregates. See also Figure 1.

**Figure S2.**
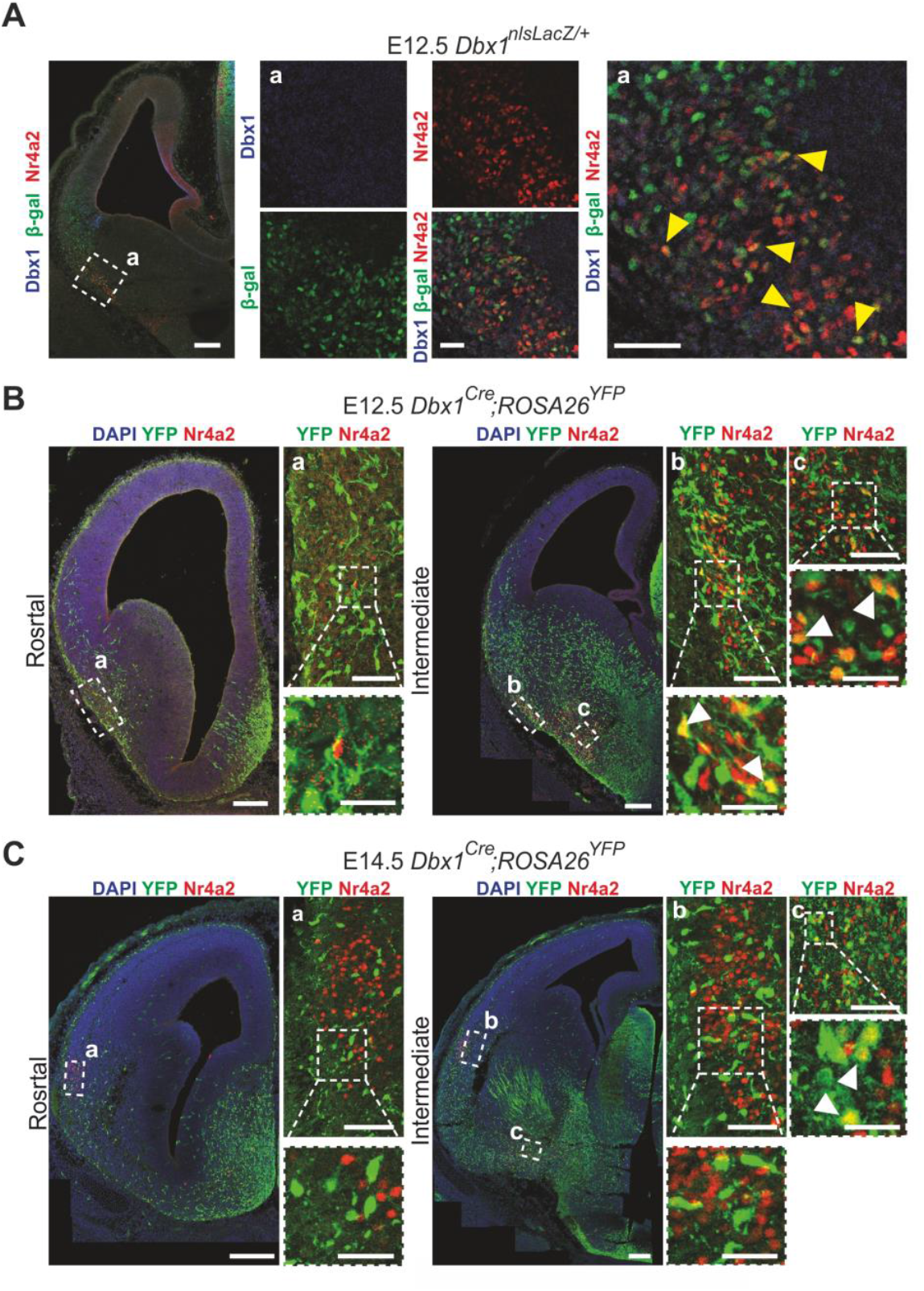
Dbx1-derived cells express Nr4a2 in the ventral pallium. (A) Confocal images of coronal brain sections from E12.5 Dbx1nlsLacZ/+ embryos, labeled for β-gal (green), Dbx1 (blue) and Nr4a2 (red). The dashed square (a) magnified on the right. Yellow arrows indicate β-gal+Nr4a2+ cells. Scale bars: 100 μm, 50 μm (a). (B, C) Confocal images of coronal brain sections from (B) E12.5 and (C) E14.5 Dbx1Cre;Rosa26YFP embryos labeled with YFP (green), Nr4a2 (red) and DAPI (blue). Dashed boxes (a, b, c) magnified on the right of each panel. Dashed squares highlighting selected groups of cells in panels a, b and c magnified below. White arrowheads indicate YFP+Nr4a2+ cells. Scale bars: 100 μm; 50 μm (top magnified); 25 μm (bottom magnified). See also Figure 1.

**Figure S3.**
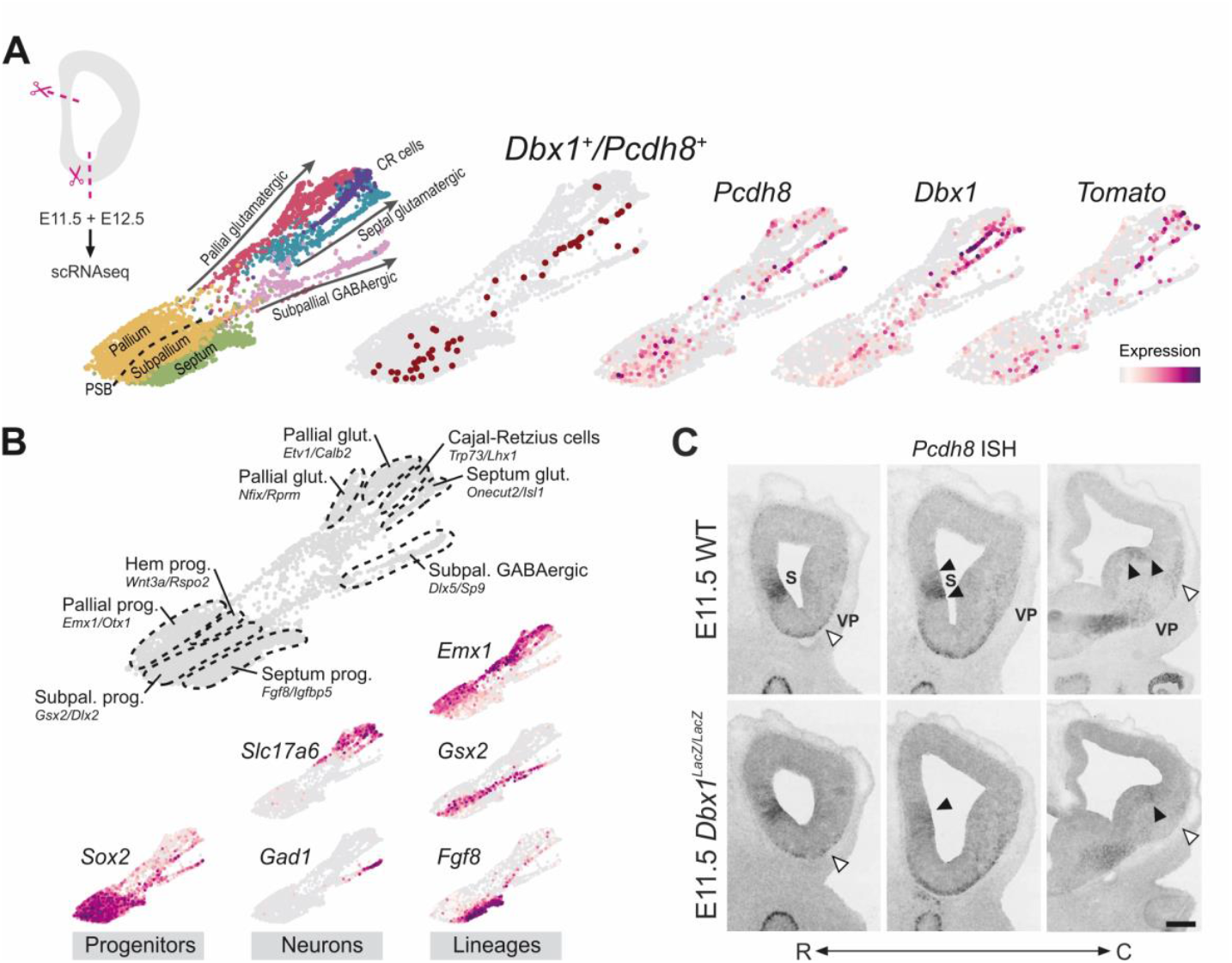
Distinct lineages and differentiation trajectories are characterized around the septum. (A) Upper left: Schematic representation of the dissection used for scRNAseq profiling. Top: SPRING dimensionality reduction of the septum dataset, with cells colored by cell type and origin. Arrows indicate the main differentiation trajectories, and the dashed line marks the pallial-subpallial boundary (PSB). CR, Cajal-Retzius. Cells co-expressing Dbx1 and Pcdh8 are show in dark red (top right). Bottom: Expression levels of Pcdh8, Dbx1 and Tomato transcripts. (B) Top: SPRING representation of the septum dataset showing the distinct progenitor and neuronal populations that were identified, with selected marker genes indicated for each population. Bottom: Expression level of selected genes in marked differentiation trajectories. (C) Bright-field images of in situ hybridization (ISH) for Pcdh8 along the rostro-caudal (R-C) axis (double arrow line) in coronal brain sections of E11.5 wild-type (WT) and Dbx1LacZ/LacZ (Dbx1 KO) embryos. White and black arrowheads indicate changes of Pcdh8 expression related to the organization of cells in the ventral postmitotic compartment, and the localization of septum (S) and ventral pallium (VP) progenitor domains, respectively. Scale bar: 100 μm. See also Figure 2.

**Figure S4.**
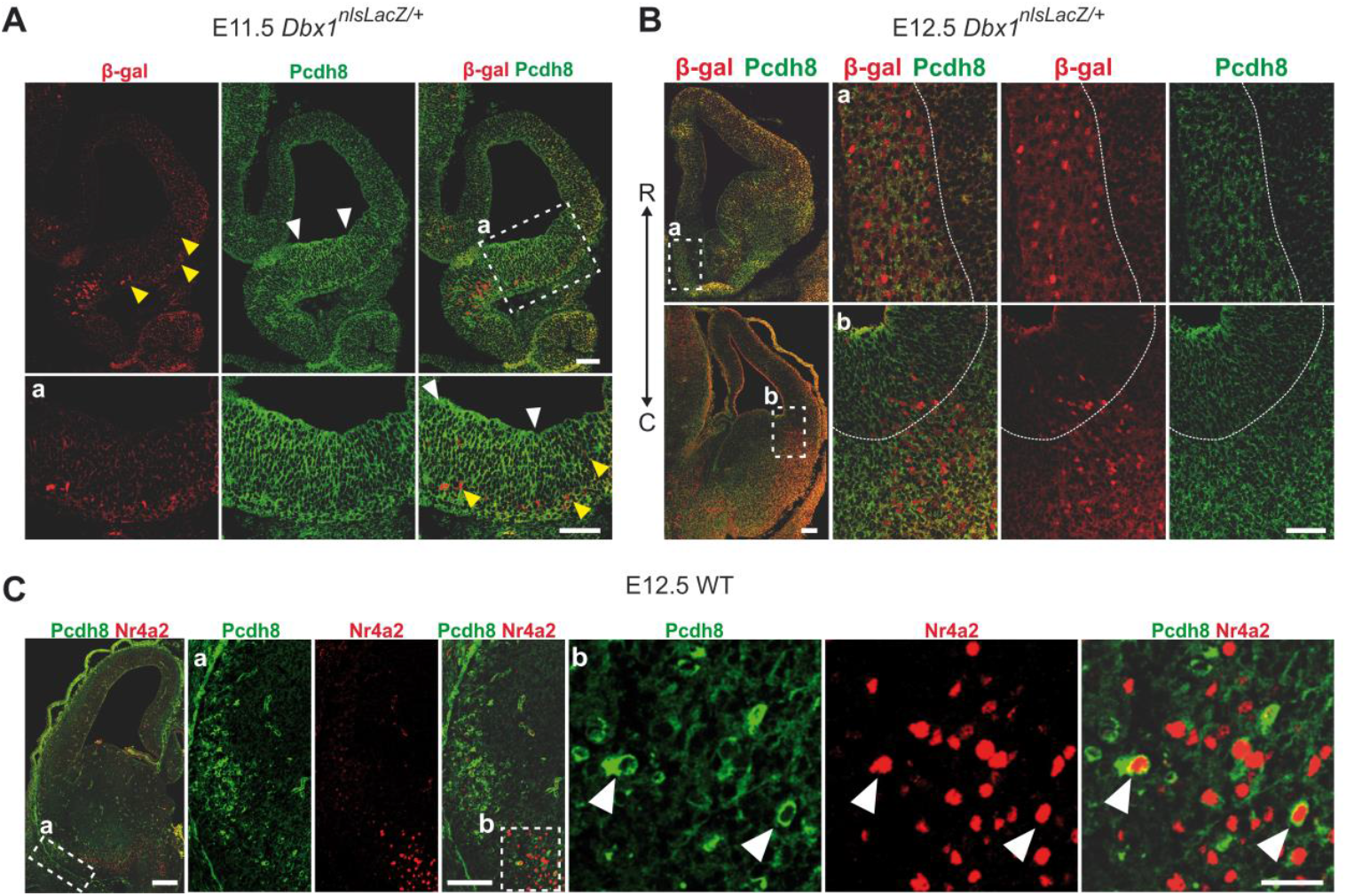
Pcdh8 and Dbx1 display interrelated expression patterns in the developing telencephalon. (A, B) Confocal images of β-gal (red) and Pcdh8 (green) co-labeling in coronal brain sections of (A) E11.5 and (B) E12.5 Dbx1nlsLacZ/+ embryos. Yellow arrowheads indicate β-gal+Pcdh8+ cells, while white arrowheads mark Pcdh8+ areas. Dashed boxes (a, b) magnified below the corresponding low-magnification images. In (B), dashed lines outline the borders of stronger Pcdh8 staining and the double arrow line indicates the rostro-caudal (R-C) axis. Scale bars: 200 μm, 100 μm (magnified). (C) Confocal images of coronal brain sections from E12.5 wild-type (WT) embryos co-labeled with Pcdh8 (green) and Nr4a2 (red). Dashed boxes (a, b) magnified on the right of the corresponding image. White arrowheads indicate Pcdh8+Nr4a2+ cells. Scale bars: 100 μm, 50 μm (a), 25 μm (b). See also Figure 2.

**Figure S5.**
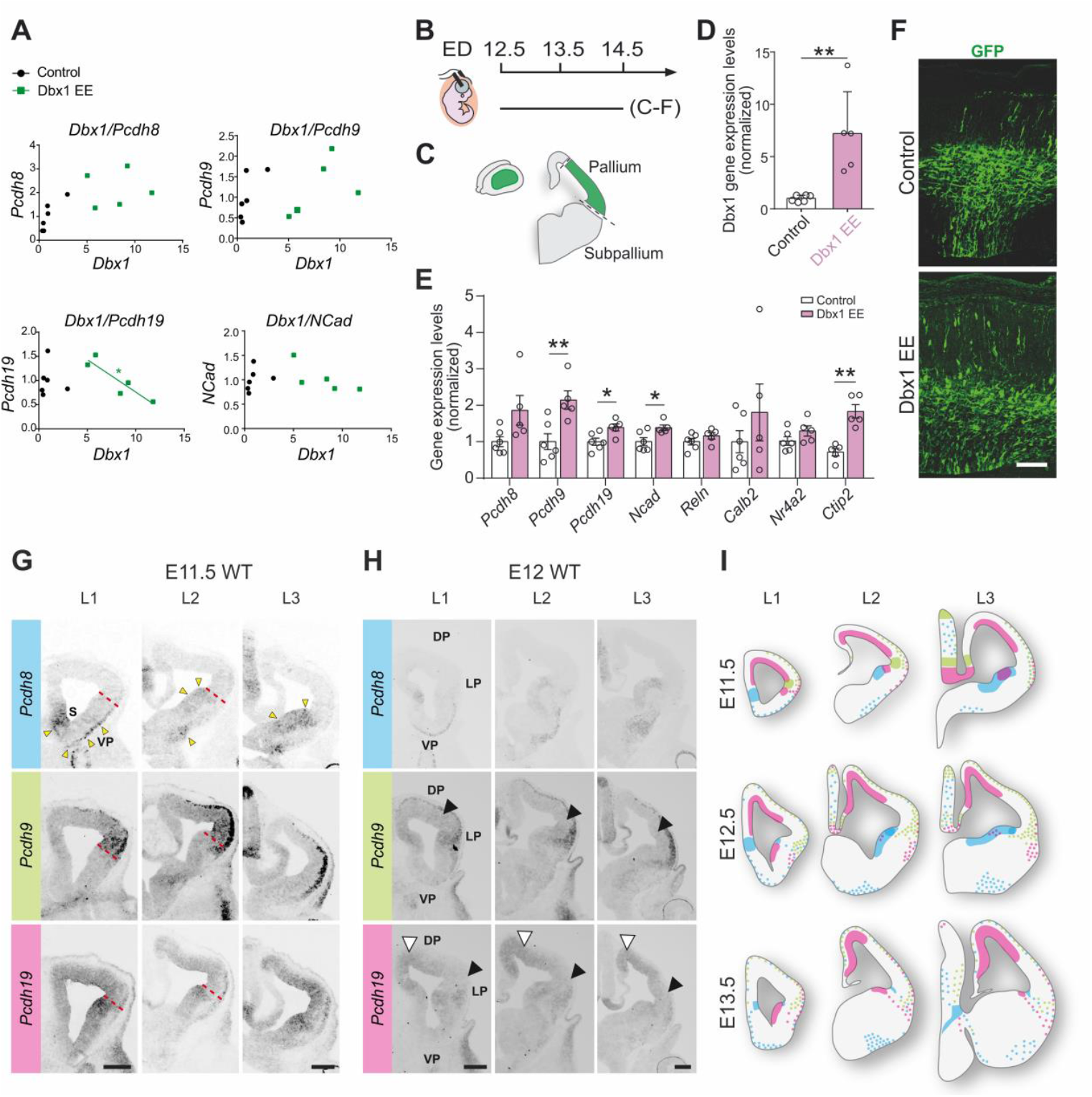
Dbx1 EE at E12.5 induces expression of different sets of genes in comparison to IUE at E11.5. (A) qPCR analysis of gene expression upon IUE (E11.5-E13.5) with either a control vector or Dbx1 EE, showing no correlation (*Dbx1/Pcdh8*; *Dbx1*/*Pcdh9*; *Dbx1/Ncad*) or a negative correlation (*Dbx1/Pcdh19*). (B) Schematic timeline of the IUE. ED, electroporation day. (C) Electroporated region dissection scheme. (D, E) qPCR quantification of the average expression levels of (D) *Dbx1* and (E) *Pcdh8, Pcdh9, Pcdh19, Ncad, Reln, Calb2, Nr4a2* and *Ctip2* upon IUE (E12.5-E14.5) with either a control vector or Dbx1 EE, normalized to the control. Data are mean ± SEM; circles represent values from independent electroporations (*n*=6 controls, *n*=5 Dbx1 EE). Student’s *t*-test: (D) \*\**p*=0.0014, (E) \**p*≤0.05; \*\**p*≤0.01. (F) Confocal images of GFP (green) in coronal sections of E14.5 mouse brains upon IUE at E12.5 with control vector or Dbx1 EE. Scale bar: 100 μm. (G, H) Representative bright-field images of *in situ* hybridization for *Pcdh8, Pcdh9* and *Pcdh19* in coronal brain sections of (G) E11.5 and (H) E12 *wild-type* (WT) embryos. Red dashed lines mark the pallial-subpallial boundary. Yellow and white arrowheads indicate regions of high *Pcdh8* or *Pcdh19* expression, respectively, while black arrowheads indicate progenitor regions with low *Pcdh9* and *Pcdh19* expression. S, septum; DP, dorsal pallium; LP, lateral pallium; VP, ventral pallium. Scale bars: 100 μm. (I) Schematic representation of *Pcdh* mRNA expression patterns at E11.5, E12.5 and E13.5. The following color code was used: blue for *Pcdh8*, green for *Pcdh9*, and pink for *Pcdh19*. L1, L2 and L3 correspond to rostral, intermediate and caudal section levels, respectively. See also Figure 3.

**Figure S6.**
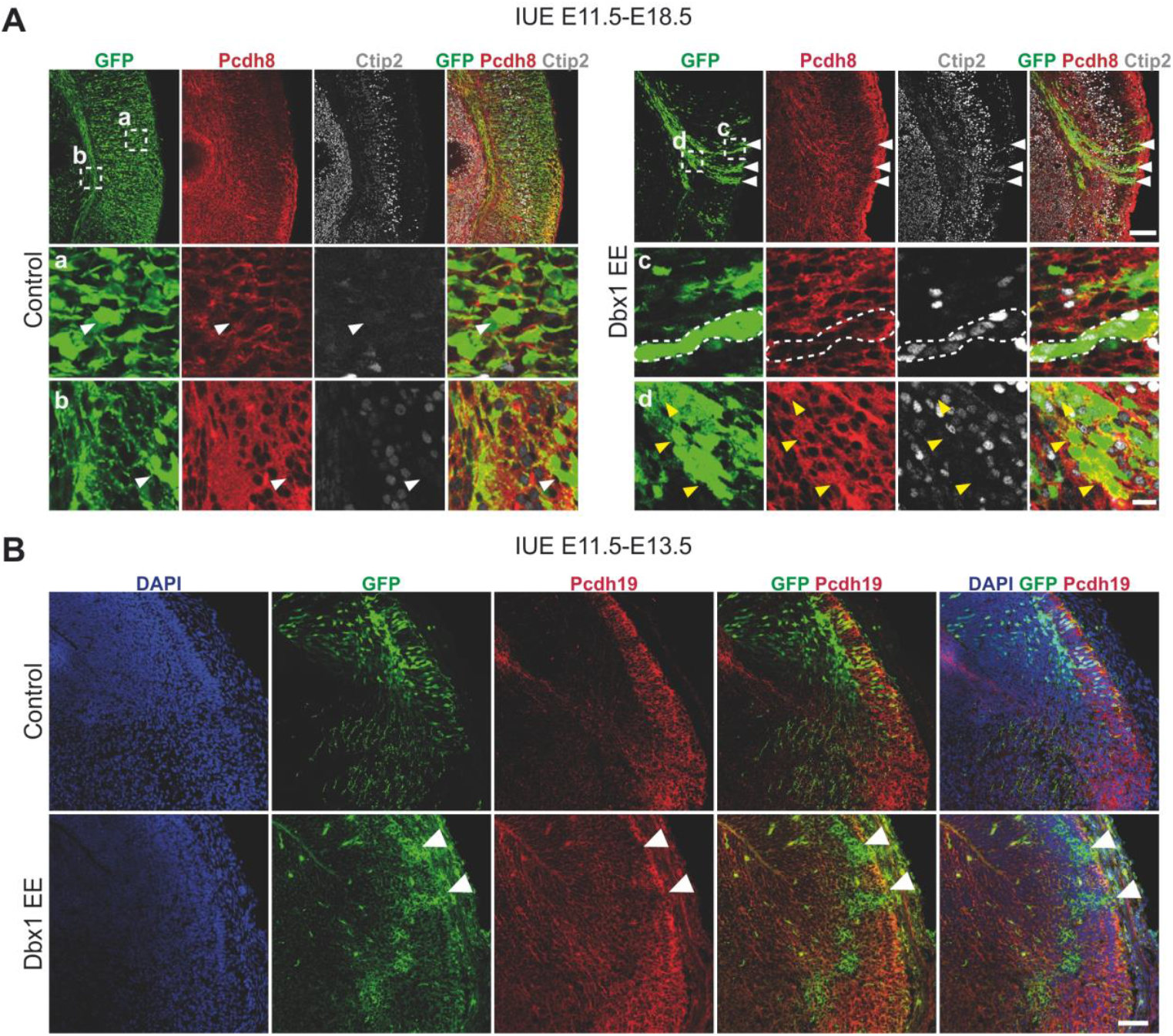
Dbx1 EE at E11.5 affects cell identity and adhesion. (A) Confocal images of GFP (green) in coronal sections of E18.5 mouse brain cortices electroporated at E11.5 with control vector or Dbx1 EE, co-labeled for Pcdh8 (red) and Ctip2 (gray). Dashed squares (a, b, c, d) magnified below. White arrowheads indicate GFP+Pcdh8-Ctip2-cells in the control and strongly adherent ‘streams’ of GFP+ cells in Dbx1 EE. The dashed area in (c) shows a group of Dbx1 EE Pcdh8lowCtip2+ cells in the cortical plate (CP), while yellow arrowheads indicate Dbx1 EE Pcdh8highCtip2-cells below the CP. Scale bars: 200 μm, 25 μm (magnified). (B) Confocal images of GFP (green) in coronal sections of E13.5 mouse brain cortices electroporated at E11.5 with control vector or Dbx1 EE, co-labeled for Pcdh19 (red) with DAPI (blue) counterstaining. White arrowheads indicate Pcdh19-aggregates induced by Dbx1 EE. Scale bar: 100 μm. See also Figure 3.

**Figure S7.**
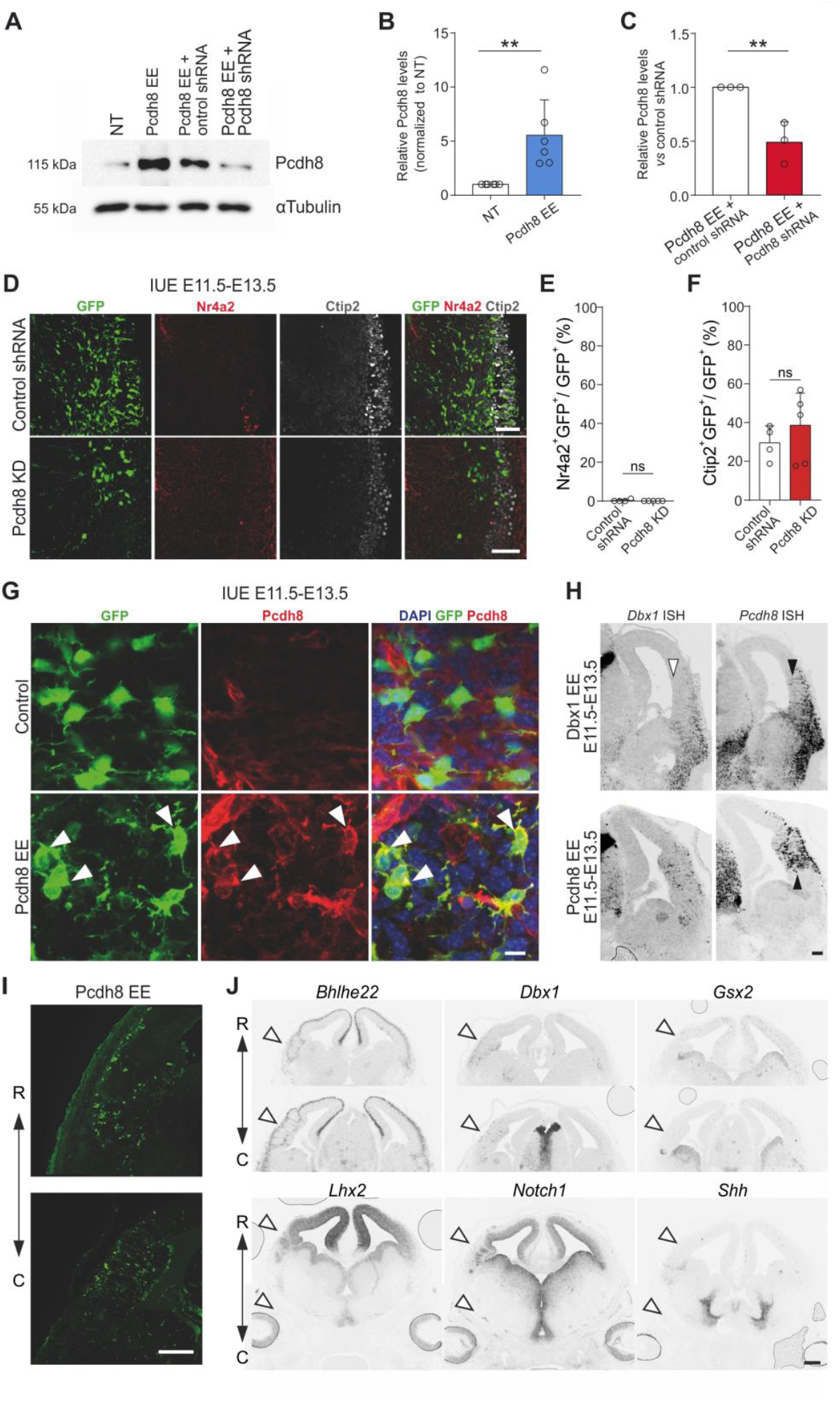
Pcdh8 KD does not affect cell identity but Pcdh8 EE alters the organization of the lateral pallium. (A) Representative immunoblotting of Pcdh8 protein levels in cell lysates from HEK293T cells 48h post-transfection under the following conditions: non-transfected (NT), Pcdh8 EE, Pcdh8 EE + control shRNA, and Pcdh8 EE + Pcdh8 shRNA. (B, C) Densitometric quantification of Pcdh8 signal, normalized to αTubulin, to assess (B) Pcdh8 EE efficacy and (C) the relative percentage of Pcdh8 knockdown (KD) (*n*=6 NT and Pcdh8 EE; *n*=3 Pcdh8 EE + control shRNA and Pcdh8 EE + Pcdh8 shRNA). Data are mean ± SEM; circles represent values from independent transfections. Student’s *t*-test: \*\**p*≤0.01. (D) Confocal images of GFP (green) in coronal sections of E13.5 mouse brains electroporated at E11.5 with a control shRNA or Pcdh8-targeted shRNA (Pcdh8 KD), co-labeled with Nr4a2 (red) and Ctip2 (gray). Scale bars: 50 μm. (E-F) Percentage of (E) Nr4a2+GFP + and (F) Ctip2+GFP+ cells among GFP+ cells. Data are mean ± SEM; circles represent values from independent electroporations (*n*=4 controls, *n*=5 Pcdh8 KD). Student’s *t*-test: ns, not significant. (G) Confocal images of GFP (green) in coronal sections of E13.5 mouse brains electroporated at E11.5 with control vector or Pcdh8 EE, colabeled with Pcdh8 (red) and DAPI (blue) counterstaining. White arrowheads indicate cells overexpressing Pcdh8. Scale bar: 50 μm. (H) Representative bright-field images of *in situ* hybridization (ISH) for *Dbx1* and *Pcdh8* in coronal sections of E13.5 mouse brains electroporated at E11.5 with Dbx1 EE or Pcdh8 EE. White arrowheads show small number of Dbx1-expressing cells in the ventricular zone, while black arrowheads show cells in the postmitotic compartment. Scale bar: 150 μm. (I) Confocal images of GFP (green) along the rostro-caudal (R-C) axis in coronal sections of E13.5 mouse brains electroporated with Pcdh8 EE. Scale bar: 200 μm. (J) Bright-field images of ISH for *Bhlhe22, Dbx1, Gxs2, Lhx2, Notch1* and *Shh* along the R-C axis in coronal sections of E13.5 mouse brains electroporated at E11.5 with Pcdh8 EE. White arrowheads indicate changes in expression along the dorso-ventral axis. Scale bar: 250 μm. See also Figures 4 and 5.

**Figure S8.**
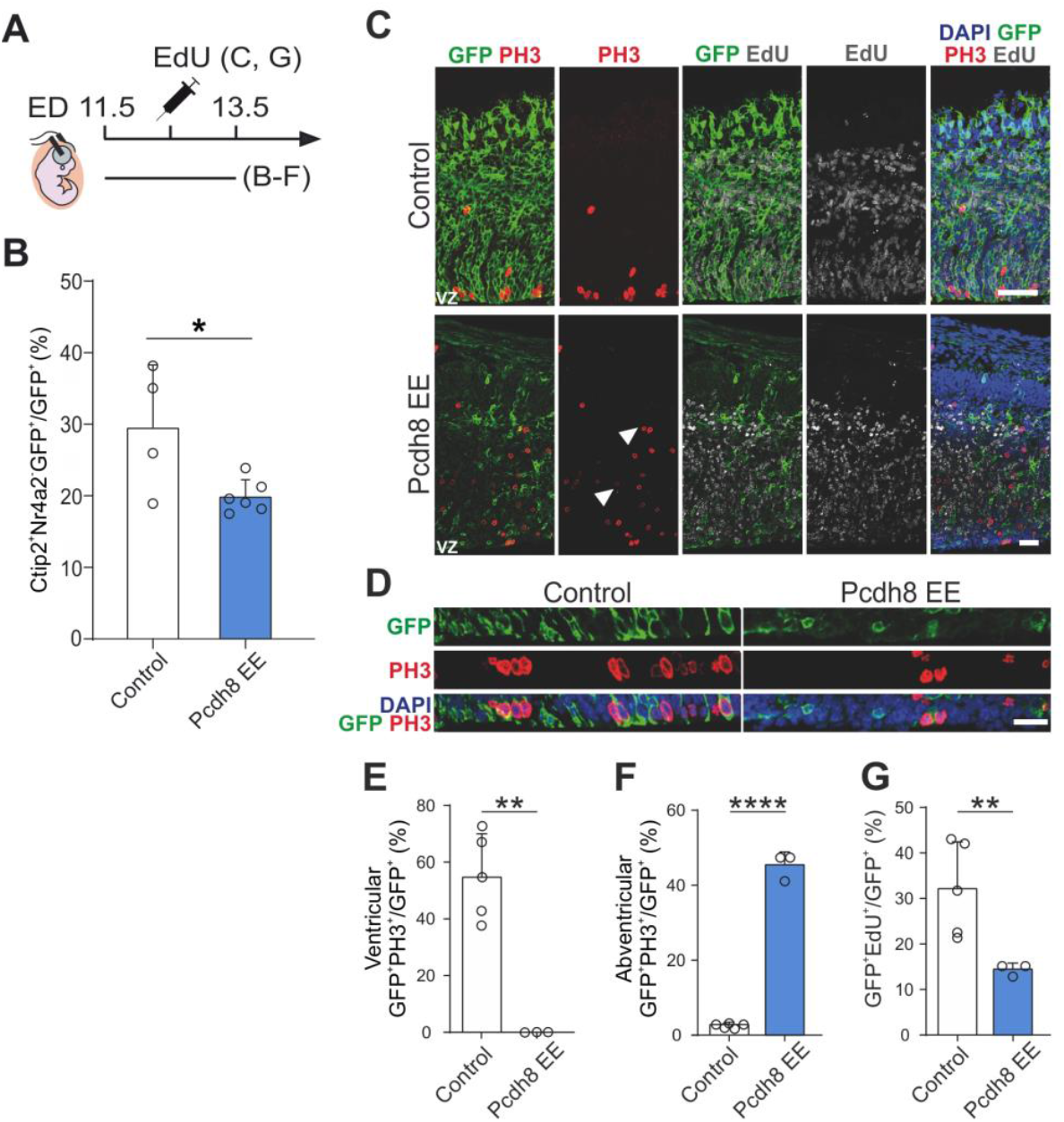
Pcdh8 EE alters the cell cycle exit. (A) Schematic timeline of the IUE. ED, electroporation day. (B) Percentage of Ctip2+Nr4a2-GFP+ cells among GFP+ cells. Data are mean ± SEM; circles represent values from individual sections (*n*=3 electroporations/condition, 1-2 sections/electroporation). Student’s *t*-test: \**p*=0.0304. (C) Confocal images of GFP (green) in coronal sections of E13.5 mouse brains electroporated at E11.5 with control vector or Pcdh8 EE, followed by EdU administration 24h post-electroporation. Sections are co-labeled with PH3 (red), EdU (gray) and DAPI (blue) counterstaining. White arrowheads indicate PH3+ cells outside the ventricular zone (VZ). Scale bars: 100 μm. (D) Electroporated VZ from (C) magnified, showing GFP (green), PH3 (red) and DAPI (blue) labeling. Scale bar: 50 μm. (E, F) Percentages of GFP+PH3+ cells among GFP+ cells, aligned either (E) at the VZ (ventricular) or (F) outside the VZ (abventricular). Data are mean ± SEM; circles represent values from independent electroporations (*n*=3-5 mice/condition, 2 sections/mouse). Student’s *t*-test: (E) \*\**p*=0.01, (F) \*\*\*\**p*<0.001. (G) Percentage of GFP+ EdU+ cells among GFP+ cells. Data are mean ± SEM; circles represent values from independent electroporations (*n*=3-5 mice/condition, 2 sections/mouse). Student’s *t*-test: \*\**p*=0.0087. See also Figure 6.

## MATERIAL AND METHODS

### Animals

The following mouse lines were used and maintained on a *C57BL/6J* background: *Dbx1*^*LacZ*^ (Pierani et al., 2001), *Rosa26*^*YFP*^ (Srinivas et al., 2001) and *Rosa26*^*tdTomato*^ (Madisen et al., 2010). All animals were handled in strict accordance with good animal practice as defined by the national animal welfare bodies, and all mouse work was approved by the Veterinary Services of Paris (Authorization number: 75–1454) and by the Animal Experimentation Ethical Committee Paris-Descartes(CEEA-34) (Reference: 2018012612027541).

### *In utero* electroporation

The procedure was carried out following previously published protocols (Arai et al., 2019; Szczurkowska et al., 2016). Briefly, to ensure the welfare of the used animals, 30 min before the surgery 50 µg/kg of Buprenorphine followed by 5 mg/kg Ketoprofen post-surgery subcutaneous administration were performed. *Wild-type C57BL/6J* pregnant mice at E11.5 or E12.5 were subjected to incision under anesthesia with Isoflurane (AXIENCE SAS), and the uterine horns were exposed onto 1X phosphate-buffered saline (PBS)-moistened cotton gauze. Embryos were visualized using appropriate flexible light sources through the uterus. Plasmid DNAs mixed with a filtered Fast Green dye were injected into the lateral ventricle through a glass capillary. A pair of 3 mm electrodes was applied to the embryos through the yolk sac, and a series of square-wave current pulses (25 V, 50 ms) was delivered for six times at 950 ms intervals using a pulse generator (NEPA21, NEPAGENE). Uterine horns were repositioned into the abdominal cavity, and the abdominal wall and skin were sutured. The concentration of plasmid DNAs used were between 1-3 μg/μl. Rescue experiments with Pcdh8EE and Jag1EE were conducted using a 1:1 ratio. The animals were checked in the post-surgery care room (heating pad, wet food and, if necessary, additional dose of painkiller was administrated).

### Plasmid DNA cloning

For the study we used a pCAGGS-HA-mouseDbx1-ires-NLS-EGFP vector previously published in Arai et al. (Arai et al., 2019). The Pcdh8 overexpression sequence was subcloned into the pCAGGS vector backbone using In-Fusion® HD Cloning (Takara Bio) accordingly to the manufacturer’s instructions.

Pcdh8 shRNA and scrambled shRNA sequences were subcloned into psiSTRIKECAG-ires-GFP vectors (a gift from Dr. Billuart) using In-Fusion® HD Cloning (Takara Bio) accordingly to the manufacturer’s instructions.

### Immunofluorescence

Mice were sacrificed by cervical dislocation and embryos were collected and fixed at 4°C for 2h (E11.5-E12.5), 2.5h (E13.5), 3h (E14.5), 4h (E18.5) in 4% paraformaldehyde (PFA)/PBS, washed in PBS 3x1h at 4°C, cryoprotected in 30% sucrose/PBS overnight at 4°C, and embedded in Tissue-Tek O.C.T compound (Sakura Finetek, Europe). Coronal cryosections of mouse embryos (20 µm: E11.5-E14.5; 25 µm: E18.5) were blocked with 0.2% Triton X-100 in 10% horse serum/1X PBS for 30 min. For Pcdh8 immunostaining, before blocking, sections were incubated for 10 min at 95°C in antigen retrieval solution (10mM sodium citrate, pH 6.0) and cooled down for 20 min at room temperature (RT). Sections were subsequently incubated overnight at 4°C with primary antibodies, followed by incubation with fluorescently labeled secondary antibodies and DAPI in 0.2% Triton X-100, 1% horse serum, 1X PBS buffer for 45 min at RT. Sections were mounted with Vectashield mounting medium. Primary antibodies used were: Calb 1:1000; Ctip2 1:2000; GFP 1:2000; Dbx1 1:10000; Delta1 1:100; Jag1 1:500; Ncad 1:400; Nurr1/Nr4a2 1:200; Pax6 1:1000; Pcdh8 1:700; Pcdh19 1:500; PH3 1:500; Reln 1:1000; Tle4 1:500. Secondary antibodies used were: Alexa 488 1:700; Cy3 1:700; Cy5 1:500

### cDNA synthesis and qPCR

RNA samples from electroporated tissue were retro-transcribed to cDNA using the RevertAid First First Strand cDNA Synthesis Ki, according to the supplier’s instructions. For qPCR analysis of gene expression in each electroporated region we used 500 ng of RNA. The genomic DNA was removed by incubation for 30 min in 37°C with DNase I, RNase-free. All qPCR reactions were performed on an Eppendorf machine in 20 μl reactio volume of Go Tag qPCR Master Mix (1× SYBR®Green PCR Master Mix and 200 nM of each primer). The following thermal program was applied: denaturation for 2 min at 95°C followed by 40 amplification cycles of 15 sec denaturing step (95°C) and 1 min annealing–extension step (60°C). Afterwards, to verify that results are coming from the single transcript the automatic program for melting curve analysis was performed using standard machine settings.

### *In situ* and fluorescent *in situ* hybridization

*In situ* hybridization was performed as previously described (Griveau et al., 2010). For each gene of interest, a DNA fragment (typically 500-800 bp) was amplified by PCRs from an embryonic brain cDNA library using Phusion polymerase (Thermo) and primers indicated in Table 1. The promoter sequence of the T7 RNA polymerase (GGTAATACGACTCACTATAGGG) was added 5′ of the reverse primers. Antisense RNA probes were labeled using a DIG-RNA labeling mix (Roche). Alternatively, for mouse *Gsx2, Lhx2* (gift from Dr S. Retaux), *Notch1* (gift from Prof U. Lendahl) and *Shh* (gift from Dr S. Garel), a plasmid containing part of the cDNA was linearized by restriction enzyme digestion and subsequently submitted to reverse transcription. Sections were mounted with Mowiol 4-88. For fluorescent *in situ* hybridization (FISH), antisense RNA probes were labeled using either the DIG-RNA labeling mix (Roche) or the Fluorescein labeling mix (Roche). Fluorescent detection was performed using Anti-FITC-POD (Roche) or Anti-DIG-POD (Roche), followed by signal amplification with Alexa Fluor™ 488 Tyramide SuperBoost™ (Invitrogen) or Cyanine 3 Tyramide (Biotechne), respectively.

**Table 1.**
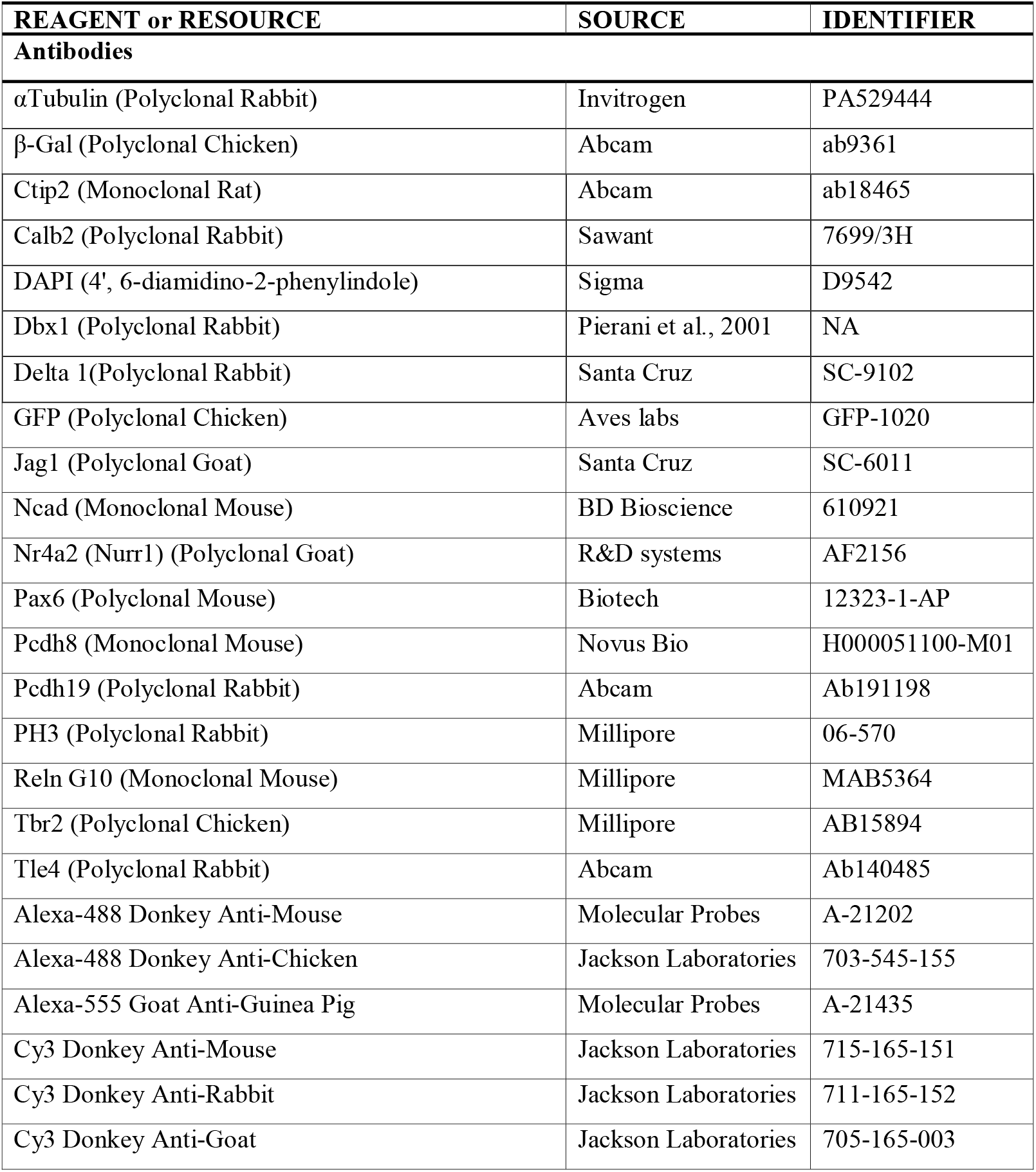
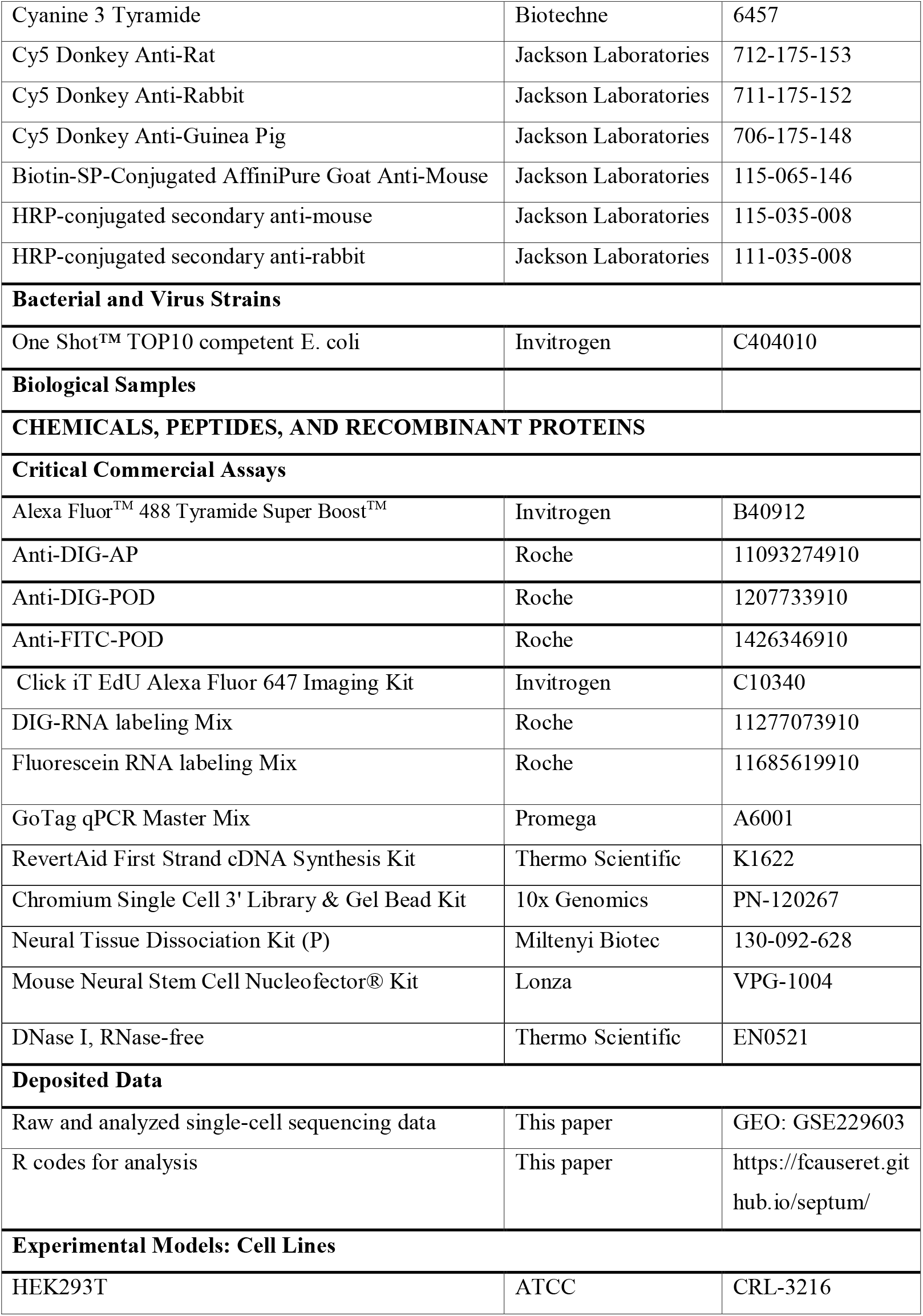
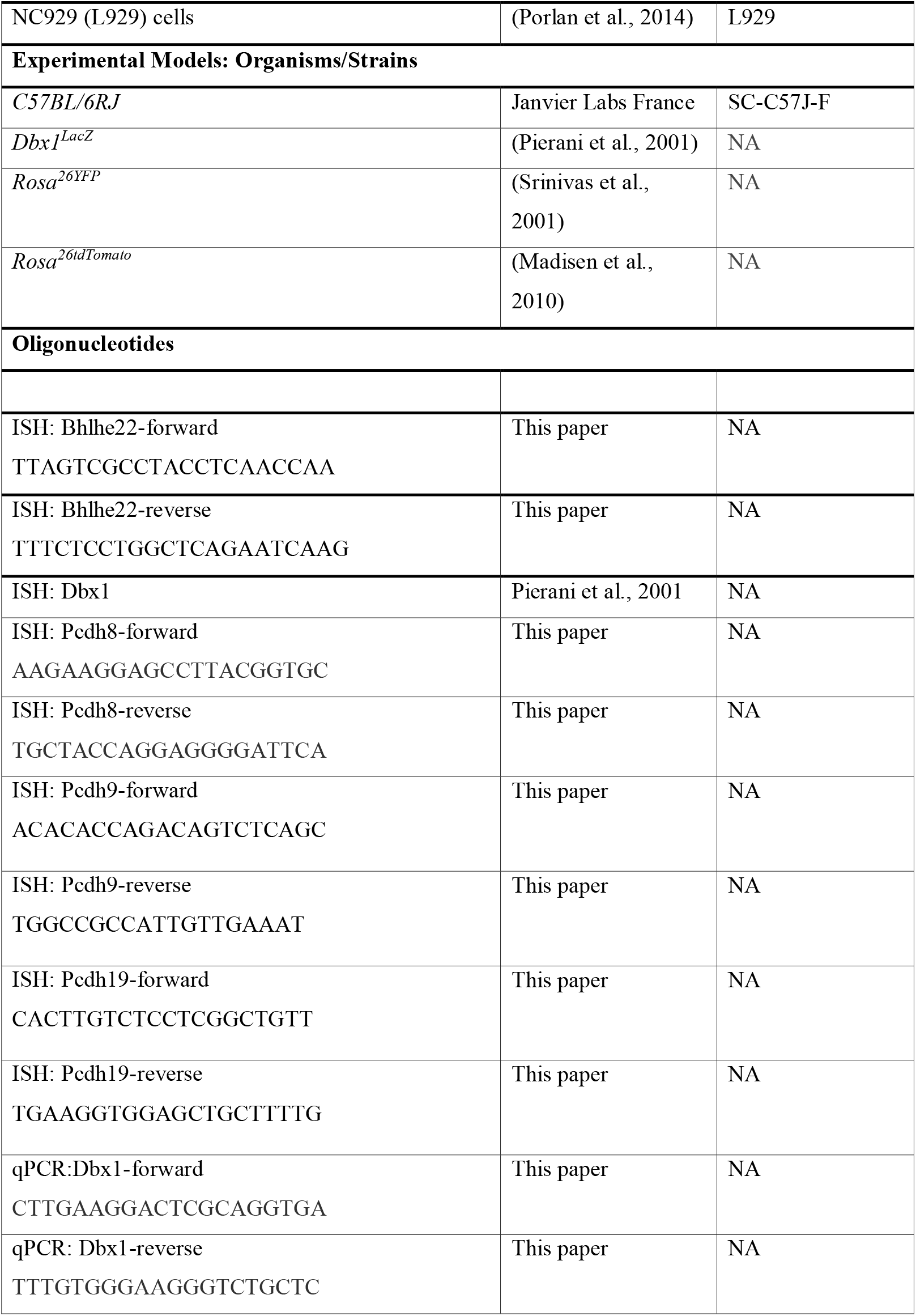
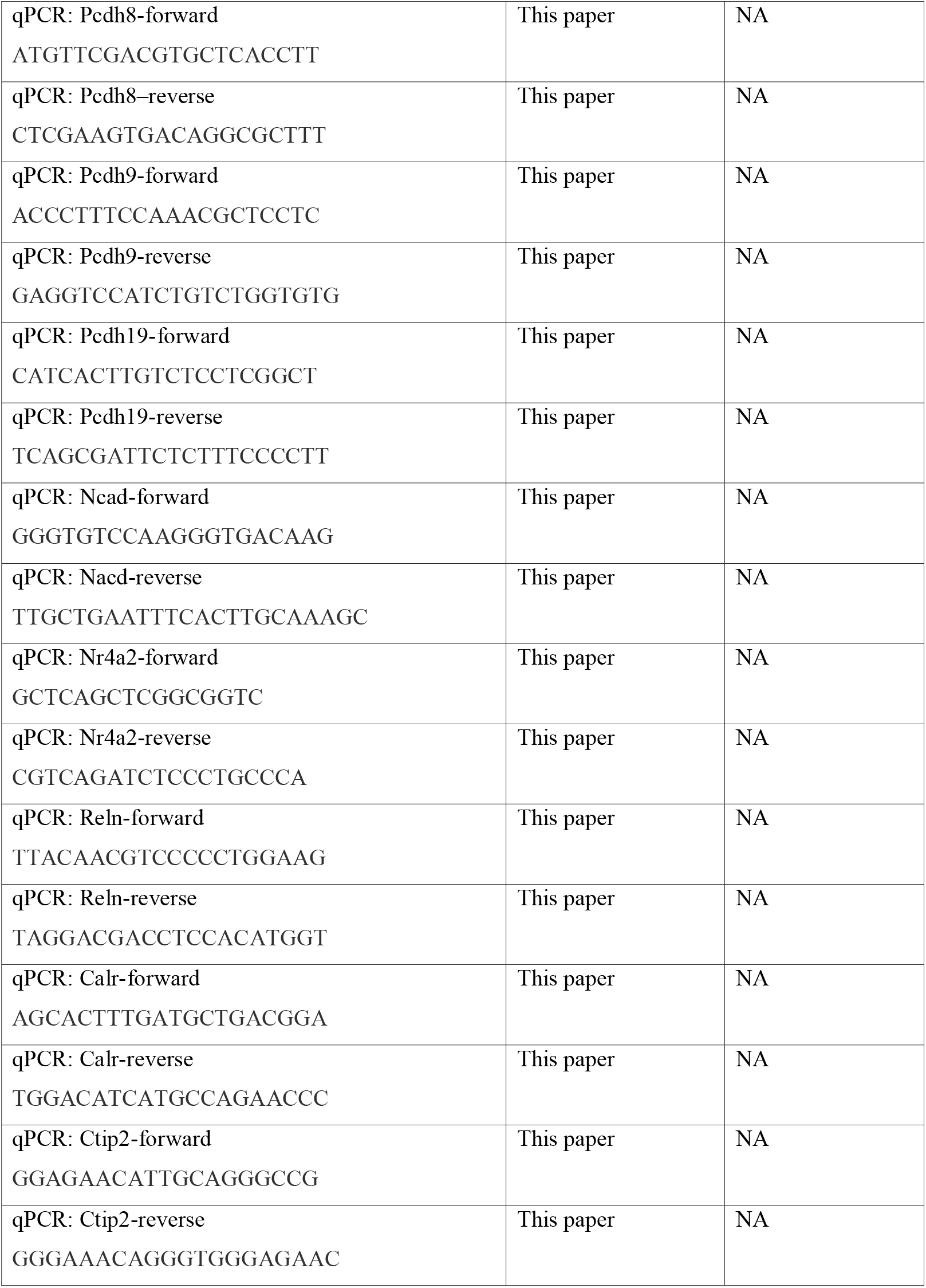
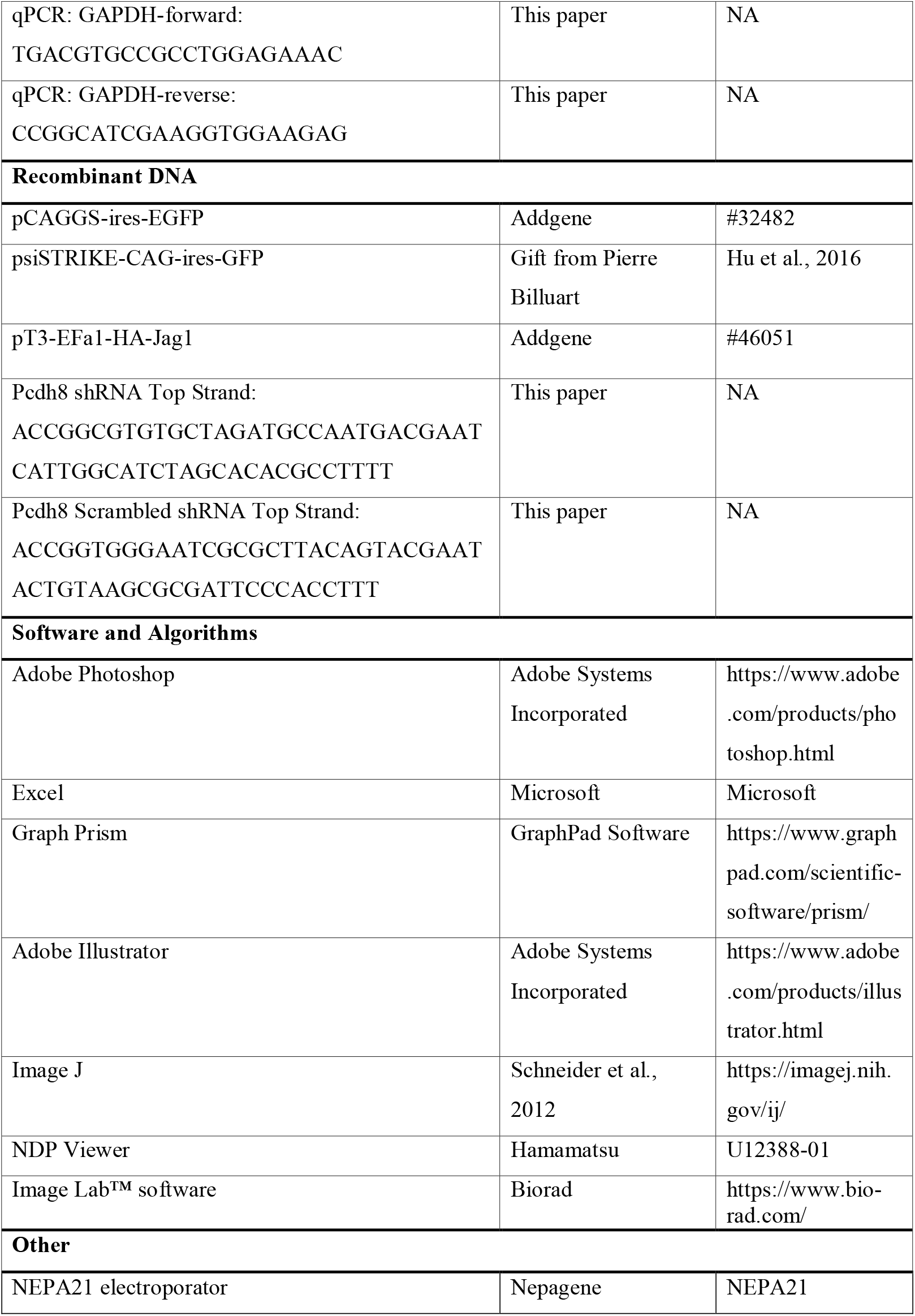
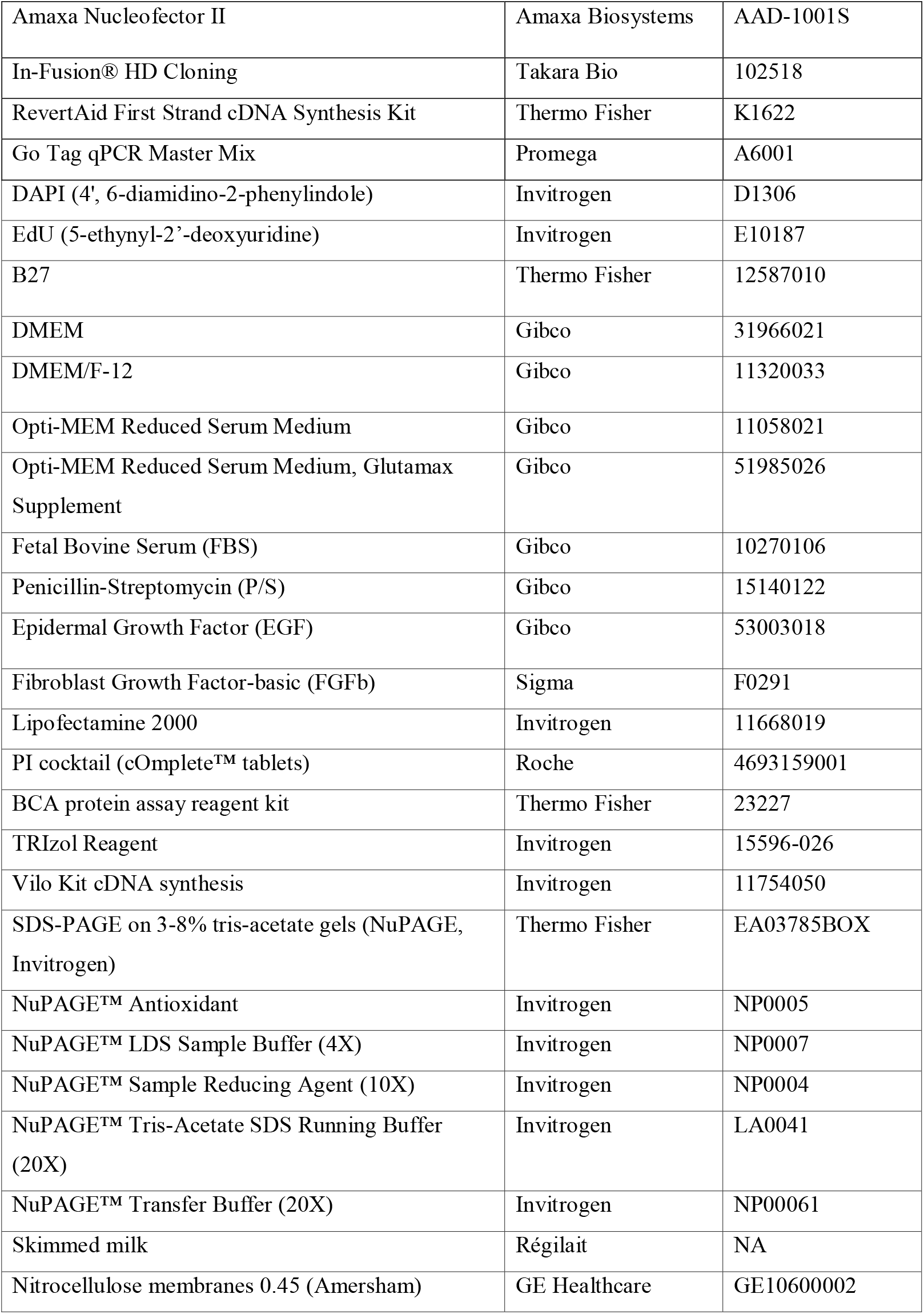
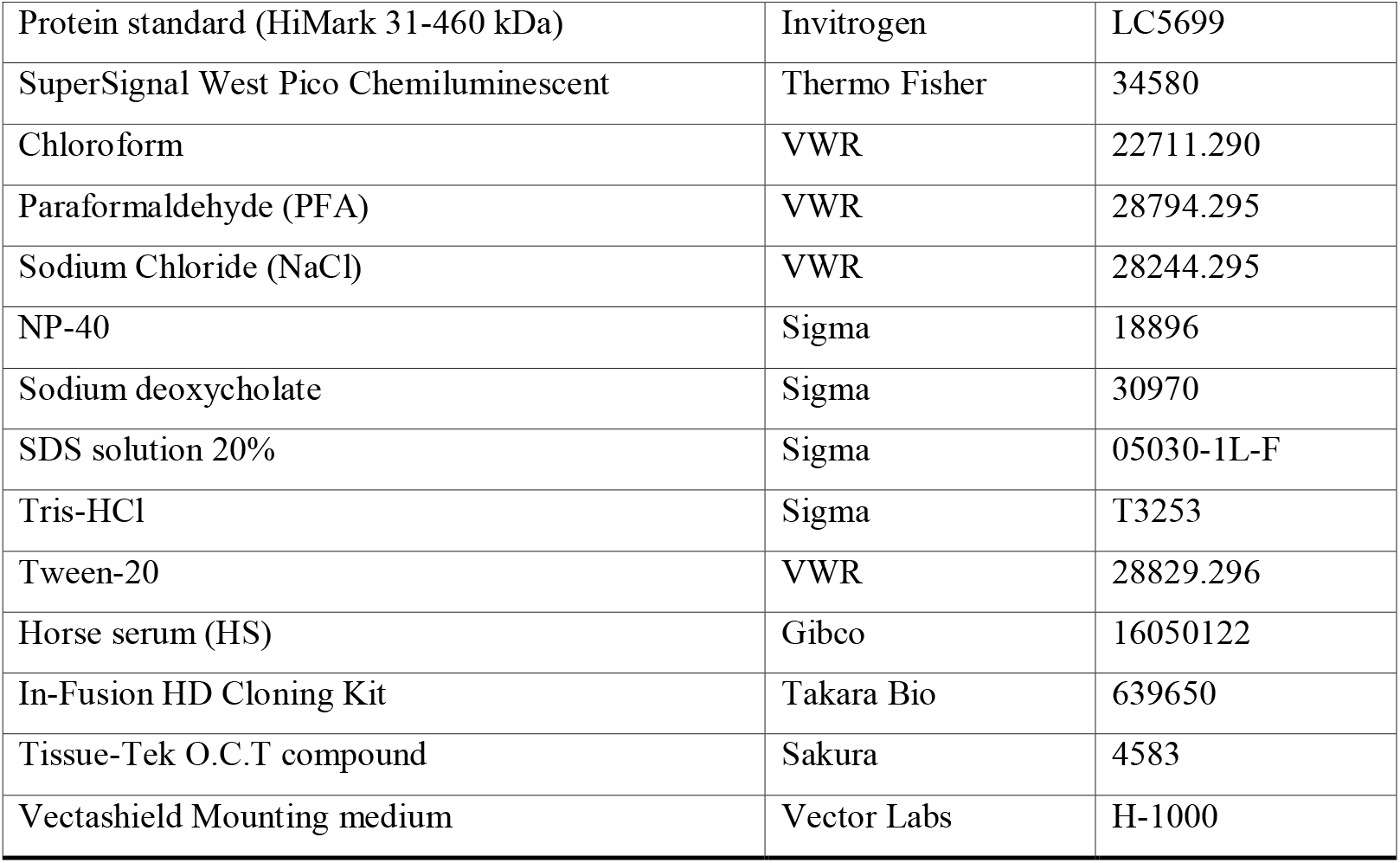
List of reagents used in the study.

### Image acquisition and quantifications

*In situ* hybridization images were obtained using a Hamamatsu Nanozoomer 2.0 slide scanner (Hamamatsu Nanozoomer 2.0HT) with a 20X objective. The scans were verified using NDP Viewers and imported into Adobe Photoshop for figure adjustments. All fluorescence images and tile scan images were acquired using a Leica TCS SP8 SMD confocal microscope equipped with a 40X objective, analyzed with ImageJ and imported into Adobe Photoshop for figure adjustments. Quantifications were performed by counting the proportion of GFP^+^ cells among the selected marker positive population using the ImageJ software. For consistency, the data shown in Figures 1E, 1F, S1D, 4D, 4E and 4F for control and Dbx1 EE originate from the same set of experiments. These data are included in multiple figures to enable comparisons across different experimental conditions while minimizing the number of animals used.

### EdU pulse labeling and staining

EdU injection was carried out by intraperitoneal injection of 100 μL of 1 mg/ml EdU (Invitrogen) in PBS into pregnant females at E12.5 after electroporation at E11.5. Immunofluorescence and EdU staining was performed using Click iT EdU Alexa Fluor 647 Imaging Kit (Invitrogen).

### Cell culture, transfection, sample preparation and adhesion assay

Human embryonic kidney (HEK) 293T cells were grown in Dulbecco’s modified Eagle’s medium (DMEM; Gibco) containing 10% fetal bovine serum (FBS) and 1% penicillin-streptomycin (P/S; Gibco) and maintained at 37°C in a 5% CO_2_ atmosphere. Cells were seeded onto 6-multiwell plates at a density of 62500 cells/cm^2^. Twenty-four hours later, transfection with 2 μg of pCAGGS constructs containing control GFP or Pcdh8, and with 4 μg of Pcdh8 scrambled shRNA or Pcdh8 shRNA was performed for 4h using Lipofectamine 2000 (Invitrogen) according to the manufacturer’s instructions. Non-transfected cells (treated with transfection reagent only) were also used as control. Cells were maintained in serum-free Opti-MEM (Gibco) supplemented with 1% P/S after transfection and cultured for an additional 40h. Cell lysates were collected using RIPA buffer (50 mM Tris-HCl pH 7.5, 150 mM NaCl, 1% NP-40, 0.1% SDS, 0.5% sodium deoxycholate) supplemented with 2 mM EDTA and protease inhibitors (cOmplete™ EDTA-free tablets, Roche). Samples were then left in a rotator for 30 min at 4°C to better dissolve the proteins and centrifuged at max speed for 10 min at 4°C in order to pellet out crude undissolved fractions. The supernatant was stored at −20°C until use. Protein quantification was determined using the bicinchoninic acid (BCA) protein assay reagent kit (Pierce™, ThermoFisher Scientific, USA) using bovine serum albumin (BSA) as standard.

L929 cells were plated and maintained in Dulbecco’s Modified Eagle Medium (DMEM) supplemented with 1% L-glutamine and 10% fetal bovine serum (FBS). Cultures were incubated at 37°C in a 5% CO_2_ humidified incubator.

Adult subependymal zone (SEZ) neural stem cell (NSC) cultures were obtained from 2–3-month-old mice, following the protocol described by (Belenguer et al., 2016). NSCs were cultured in DMEM/F-12 (Gibco) medium supplemented with 20 ng/mL epidermal growth factor (EGF, Gibco), 10 ng/mL basic fibroblast growth factor (bFGF, Sigma), and 1× B27 supplement (Thermo Fisher). Cultures were maintained for 7–10 days at 37°C in a 5% CO_2_ humidified incubator.

For adhesion experiments, L929 cells overexpressing N-cadherin were seeded onto glass coverslips and allowed to reach confluence over 24–48 hours. Passage 2 NSCs were freshly dissociated from the SEZ of three adult *wild-type* (WT) mice (2–3 months old) and transfected with either a Control or Dbx1EE vector using the NSC Nucleofection Kit (Lonza) and Amaxa Nucleofector II (Amaxa), following the manufacturer’s instructions. After 24 hours, transfected cells were analyzed via fluorescence-activated cell sorting (BD LSRFortessa™ Cell Analyzer) to determine the percentage of GFP^+^ cells. Subsequently, NSCs were seeded at a density of 1.25 × 10^4^ cells/cm^2^ onto the pre-formed L929 monolayers. After 45 minutes of incubation, cultures were thoroughly washed to remove unattached cells, followed by fixation and analysis.

### Western blot analysis

Proteins samples were added to 1/4 volume of 4X LDS sample buffer (141 mM Tris, 106 mM Tris-HCl, 2% LDS, 10% glycerol, 0.51 mM EDTA, 0.22 mM SERVA Blue G, 0.175 mM phenol red, pH 8.5), to 1/10 of sample reducing agent 10X (50 mM DTT) and boiled at 95°C for 5 min. Protein samples (10 µg of lysates) were separated by SDS-PAGE on 3-8% tris-acetate gels (NuPAGE, Invitrogen) under reducing conditions at 150 V at 4°C and electro-transferred to 0.45 µm nitrocellulose membranes for 1h30-2h at 0.5 A at 4°C. After 1h blocking at RT with 5% (w/v) milk in Tris-buffered saline (50 mM Tris, 150 mM NaCl, pH 7.6) containing 0.1% Tween-20 (TBS-T), the membranes were incubated overnight at 4°C with the following primary antibodies: Pcdh8 1:1000, αTubulin 1:10000. After three washes of 10 min with TBS-T, the membranes were incubated with the appropriate HRP-conjugated secondary antibodies (1:20000) diluted in TBS-T with 5% (w/v) milk for 1h at RT. After three 10 min washes with TBS-T, blots were developed with SuperSignal West Pico Chemiluminescent Substrate (ThermoFisher Scientific) and visualized on the ChemiDoc apparatus (Bio-Rad).

Densitometry analysis was determined using Image Lab™ software (Bio-Rad). The densities of protein bands were quantified with background subtraction. The bands were normalized to αTubulin loading control. The molecular weights were determined by using an appropriate pre-stained protein standard (HiMark 31-460 kDa, Invitrogen).

### Microarray and scRNAseq analysis

*Dbx1*^*Cre*^*;Rosa26*^*YFP*^ cell sorting and microarray analysis at E12.5 were performed as previously described in (Griveau et al., 2010). scRNAseq was performed as previously described for the VP (Moreau et al., 2021). The entire telencephalic vesicle dataset was generated using three WT embryos aged E11.5 to E12 from two distinct litters. For the septum analysis, explants encompassing the septum were dissected and pooled from eight E11.5 *PGK*^*Cre*^;*Rosa26*^*YFP*^ embryos originating from two distinct litters and four E12.5 *Dbx1*^*Cre*^;*Rosa26*^*tdTomato*^ embryos from two litters. The dissected tissue was dissociated using the Neural Tissue Dissociation Kit (P) (Miltenyi Biotec) and a gentleMACS Octo Dissociator following the manufacturer’s instructions. Cell clumps and debris were removed via filtration through 30 µm cell strainers (Miltenyi Biotec) followed by two consecutive rounds of centrifugation for 3 min at 200 *g*. Pipetting through Gel Saver tips (QSP) and a final filtration using a 10 µm cell strainer (pluriSelect) allowed to obtain a single-cell suspension. Approximatively 10000 cells were used as input on the 10X Genomics Chromium controller. A Single Cell 3′ Kit v2 library was produced and sequenced on a NextSeq500 sequencer at a total depth of 476 million reads. Raw sequencing reads were processed to count matrix with Cell Ranger version 2.1.1 using default parameters and the mm10 mouse genome reference to which the sequenced of *YFP* and *tdTomato* were added. Bioinformatic analyses were performed with a R version 4.1.1. Cells were filtered based on the percentage of mitochondrial reads, removing those outside three median absolute deviation (MAD) from the median. The count matrix was library size normalized using Seurat V4.1.0 (Hao et al., 2021). Doublet removal was achieved using Scrublet (Wolock et al., 2019). In some instances, the SPRING tool (Weinreb et al., 2018) was used for dimensionality reduction.

## Statistical Analysis

All data are expressed as mean ± SEM. *P*-values < 0.05 were considered significant and set as follows: \**p*<0.05, \*\**p*<0.01, \*\*\**p*<0.001, \*\*\*\**p*<0.0001. According to the data structure, two-group comparisons were performed using two-tailed unpaired Student *t*-test or Holm-Sidak as *post-hoc* test following one-way ANOVA. Statistics and plotting were performed using GraphPad Prism 7.0 (GraphPad Software Inc, USA).

